# MXtrodes: MXene-infused bioelectronic interfaces for multiscale electrophysiology and stimulation

**DOI:** 10.1101/2021.03.01.433237

**Authors:** Nicolette Driscoll, Brian Erickson, Brendan B. Murphy, Andrew G. Richardson, Gregory Robbins, Nicholas V. Apollo, Tyler Mathis, Kanit Hantanasirisakul, Puneet Bagga, Sarah E. Gullbrand, Matthew Sergison, Ravinder Reddy, John A. Wolf, H. Isaac Chen, Timothy H. Lucas, Timothy Dillingham, Kathryn A. Davis, Yury Gogotsi, John D. Medaglia, Flavia Vitale

**Author notes:** corresponding author: Flavia Vitale –.

## Abstract

Soft bioelectronic interfaces for mapping and modulating excitable networks at high resolution and at large scale can enable paradigm-shifting diagnostics, monitoring, and treatment strategies. Yet, current technologies largely rely on materials and fabrication schemes that are expensive, do not scale, and critically limit the maximum attainable resolution and coverage. Solution processing is a cost-effective manufacturing alternative, but biocompatible conductive inks matching the performance of conventional metals are lacking. Here, we introduce MXtrodes, a novel class of soft, high-resolution, large-scale bioelectronic interfaces enabled by Ti_3_C_2_ MXene and scalable solution processing. We show that the electrochemical properties of MXtrodes exceed those of conventional materials, and do not require conductive gels when used in epidermal electronics. Furthermore, we validate MXtrodes in a number of applications ranging from mapping large scale neuromuscular networks in humans to delivering cortical microstimulation in small animal models. Finally, we demonstrate that MXtrodes are compatible with standard clinical neuroimaging modalities.

## Introduction

Recent advances in soft materials and electronics have fueled a new generation of bioelectronic technologies for medical diagnostics and therapeutics, healthcare monitoring, and wearable devices (*1–4*). Soft and flexible bioelectronic devices designed to safely and intimately interface with multiscale networks in the human body offer paradigm-shifting opportunities for monitoring brain activity, cardiac health, and muscle function, as well as for facilitating human-machine interactions, such as myoelectric control of advanced prostheses (*5–7*) or non-invasive brain-computer interfaces (*8*). Active bioelectronic devices can also implement intelligent control strategies based on electrical stimulation of excitable circuits for treating neurological diseases (*9, 10*), heart arrhythmias (*11, 12*), and inflammatory disorders (*13, 14*), as well as for rehabilitation therapies (*15–17*).

Despite significant progress in the field of soft, ultrathin, and conformable bioelectronics (*18–22*), conventional materials and fabrication processes remain largely inadequate for producing large-scale, high-density multielectrode arrays that can map and modulate activity in excitable tissues with high spatiotemporal resolution over broad areas. For example, thin-film processing has enabled advances in ultra-thin and flexible electronics; however, such processes rely on wafer-scale microfabrication conducted in a cleanroom facility, which is time consuming, expensive, and typically limits the maximum attainable area coverage to a few square centimeters (*1*). Recent work has demonstrated the possibility to overcome this size limit by implementing the fabrication process on large-diameter wafers (*19*), but microfabrication techniques remain difficult to scale and are still not cost-effective. Contributing to the manufacturing challenges is the fact that flexible polymeric encapsulation materials require low-temperature processing below their glass transition points and can also be incompatible with certain chemicals and solvents commonly used in microfabrication. In addition to the processing constraints, conventional bioelectronic materials – such as gold (Au), platinum (Pt), iridium (Ir), and silver/silver-chloride (Ag/AgCl) – are both costly and intrinsically limited in their electronic, mechanical, and chemical compatibility with biological tissues. For example, they present challenges in achieving low electrode-tissue interface impedance, which is essential for high-fidelity recording and for safe, effective delivery of electrical stimulation. As a result, in applications such as epidermal electronics, conventional Ag/AgCl electrodes typically require a conductive gel at the skin interface to lower the impedance enough to acquire good quality signal. The conductive gels pose a host of issues including skin irritation and impedance instability as they dry out (*23, 24*). Metal-based devices are also inadequate for coupling with clinical imaging modalities such as magnetic resonance imaging (MRI), often causing artifacts in the images even if they are considered MR safe. To realize the next generation of soft, large-scale, high-density bioelectronic interfaces for human applications, innovation in both materials and fabrication is needed.

Solution processing techniques, such as inkjet or screen printing, offer a scalable and low-cost route for fabricating large-area multielectrode arrays unencumbered by the size constraints of wafer-level methods. Despite these advantages, widespread adoption of such techniques has so far been limited by the lack of conductive inks that can produce electrodes with biocompatibility, flexibility, electronic conductivity, and electrochemical properties comparable to those made with conventional materials. High conductivity can be achieved by printing ink formulations based on metal nanoparticles, however, they typically require high annealing temperatures which are not compatible with soft polymeric substrates (*25, 26*). Furthermore, metal inks raise concerns about potential toxicity (*27*). Conducting polymer-based inks, notably those using poly(3,4-ethylenedioxythiophene) doped with poly(styrene sulfonate) (PEDOT:PSS), can be processed at ambient temperatures to generate flexible, biocompatible films, but their conductivity is generally significantly lower than metal films (*28, 29*). Finally, graphene oxide (GO) inks require an additional reduction step to be converted to the conductive reduced graphene oxide (rGO) form, which poses materials and safety issues due to the high temperatures and toxic chemicals employed in the reduction process (*30*). Furthermore, the conductivity of rGO is still far inferior to that of pristine graphene (*31*) and PEDOT:PSS films.

Transition metal carbides, nitrides, and carbonitrides (MXenes) have emerged as a new class of two-dimensional (2D) nanomaterials that enable low-cost, additive-free, solution processing and can produce biocompatible films with metallic conductivity. MXenes are 2D flakes ~1 nm in thickness and up to 10s of μm in lateral size which contain abundant surface-terminating functional groups, including hydroxyl (-OH), oxygen (-O), or fluorine (-F), which make them hydrophilic and also allow for versatile surface modification and functionalization. The unique hydrophilic nature of MXenes enables a wide range of safe, high-throughput, and scalable processing methods using simple water-based inks, including spray (*32*), spin (*33*), and dip coating (*34*), direct writing (*35*), and inkjet printing (*35, 36*). Of the large variety of MXenes that have been synthesized to date, Ti_3_C_2_ (the chemical formula is often written as Ti_3_C_2_T_x_ to account for the surface terminations, T_x_) has been the most widely studied and optimized MXene. It is made of Earth-abundant elements and no cytotoxicity has been observed in previous studies (*37*). In the past few years, Ti_3_C_2_ has attracted great attention for a number of biomedical applications (*38*), including cancer theranostics (*39*), hemodialysis (*40*), and wearable mechanical sensors (*41, 42*). Our group has pioneered the use of Ti_3_C_2_ for implantable neural microelectrodes (*43, 44*) and skin-conformable, thin-film wearable sensors (*45*). Ti_3_C_2_ has metallic behavior exhibiting electronic conductivity of up to 15,000 – 20,000 S/cm, higher than all other solution-processed 2D materials (*46, 47*). Additionally, MXenes are produced using a topdown selective etching procedure which is highly scalable compared to the bottom-up techniques such as physical and chemical vapor deposition required to synthesize many other nanomaterials (*48*).

In this work, we demonstrate a new class of flexible, multichannel, high-density bioelectronic interfaces – which we have named “MXtrodes” – that are capable of both high-fidelity recording and effective stimulation of neural and neuromuscular circuits at multiple scales. The key value of the work presented here rests on a series of advances: first, we leverage the excellent processability of Ti_3_C_2_ MXene to develop a rapid, low-cost, and highly scalable method for fabricating multichannel electrode arrays of arbitrary size and geometry. This process is conducive to industrial manufacturing, and paves the way for translating MXene bioelectronics into clinical and consumer markets. Second, we report the first comprehensive study of the electrochemical properties of Ti_3_C_2_ relevant for recording and stimulation of bioelectric circuits. We show that the electrochemical behavior of Ti_3_C_2_ is comparable, and in many aspects superior to, conventional bioelectronic materials, especially in the context of delivering charge for safe and effective electrical stimulation. Third, we demonstrate the utility of MXtrodes for mapping and modulating excitable networks at scales ranging from large neural and muscular circuits in humans to small animal models. In particular, we show gel-free multichannel arrays for large-scale human epidermal recording with electrode-skin interface impedance and recording quality comparable to larger, commercially available pre-gelled Ag/AgCl single contact electrodes. Furthermore, we demonstrate the ability of MXtrodes to finely map clinically relevant neural and neuromuscular activation patterns at high spatial and temporal resolution – which is elusive for conventional epidermal sensors – and to deliver effective electrical stimulation. Fourth, we experimentally characterize the compatibility of Ti_3_C_2_ with clinical imaging modalities and we demonstrate that MXtrodes minimally interact with magnetic fields and X-rays, resulting in artifact-free high-field MRI and computed tomography (CT) imaging. This finding opens up new and exciting opportunities for future research and clinical paradigms combining high temporal resolution electrophysiology with advanced functional imaging. Propelled by the unique properties of Ti_3_C_2_ and the high-throughput, scalable, and cost-effective fabrication process developed here, MXtrodes show great promise for numerous applications in healthcare, research, and wearable electronics.

## Results

### Rapid, low-cost manufacturing of MXtrodes

We developed a simple method for producing the MXtrode arrays involving 1) laser-patterning a porous absorbent substrate, 2) infusing it with a water-based Ti_3_C_2_ MXene ink, and 3) encapsulating the resulting conductive composite in flexible elastomeric films. For the various applications demonstrated in this work, we fabricated two types of electrodes by slightly varying the same basic process: 1) flat or planar electrodes for gel-free epidermal sensing and for epicortical brain recording and stimulation, and 2) 3D “mini-pillar” electrodes for gel-free electroencephalography (EEG). The versatility of our fabrication process allows addressing application-specific requirements and customizing the electrodes to the structures of interest: while for epidermal and cortical recording flat planar electrodes achieve adequate tissue coupling, gel-free EEG requires 3D components to overcome the hair barrier and make contact with the scalp. The fabrication process, with both variations, is depicted in Fig. 1A. Briefly, we used a CO_2_ laser to pattern a nonwoven, hydroentangled cellulose-polyester blend substrate into the desired electrode array geometry. This served as a scaffold for the MXene flakes, with the laser patterning process allowing for rapid prototyping and customization of the array geometry. Next, we infused the cellulose-polyester substrate with MXene ink, which was prepared by a minimally intensive layer delamination (MILD) method (*49*) to produce a water-based MXene ink with a concentration of 30 mg/mL. The ink quickly wicked into the absorbent substrate, coating all the fibers. The ink-infused substrates were then thoroughly dried in a vacuum oven for 1 h at 70°C and 60 mmHg. The resulting structure was a rough, macro-porous conductive composite, with MXene flakes coating the individual fibers in the textile matrix (fig. S1, A and B). For the planar MXtrode arrays, the MXene conductive composite was then encapsulated in ~1 mm-thick layers of polydimethylsiloxane (PDMS), with a thorough degassing step prior to curing that allows the PDMS to infiltrate into the conductive matrix (fig. S1C). Electrode contacts were defined by cutting through the top encapsulation layer with a biopsy punch of the desired electrode diameter and peeling up the resulting PDMS disk to expose the conductive MXene composite beneath (fig. S1D). In the planar MXtrode arrays for cortical recording and stimulation, an additional 1 μm-thick layer of parylene-C was deposited prior to opening the electrode contacts to serve as an additional barrier to moisture. To fabricate 3D MXtrode arrays for gel-free EEG recording, “mini-pillars” of MXene-infused cellulose foam were deposited onto the electrode locations prior to PDMS encapsulation. Similar to the absorbent cellulose-polyester substrate, the cellulose foam readily absorbed the MXene ink, which thoroughly coated all surfaces to form a porous conductive composite after vacuum drying (fig. S1E). It is worth noting that no adhesive was required to affix the 3D mini-pillars to the underlying laser-patterned substrate: inking the two structures simultaneously with MXene and vacuum drying them while in contact resulted in the formation of a continuous conductive network fusing the laser-patterned substrate and the cellulose foam together. The final steps of the 3D MXtrode array fabrication involved PDMS encapsulation and manual trimming of the mini-pillars to expose the conductive MXene-cellulose foam composite. The versatility, simplicity, scalability, and low cost of this process enabled parallel fabrication of MXtrodes in a variety of geometries for diverse bioelectronic applications, even within the same batch (Fig. 1, B to E, fig. S2).

**Fig. 1.**
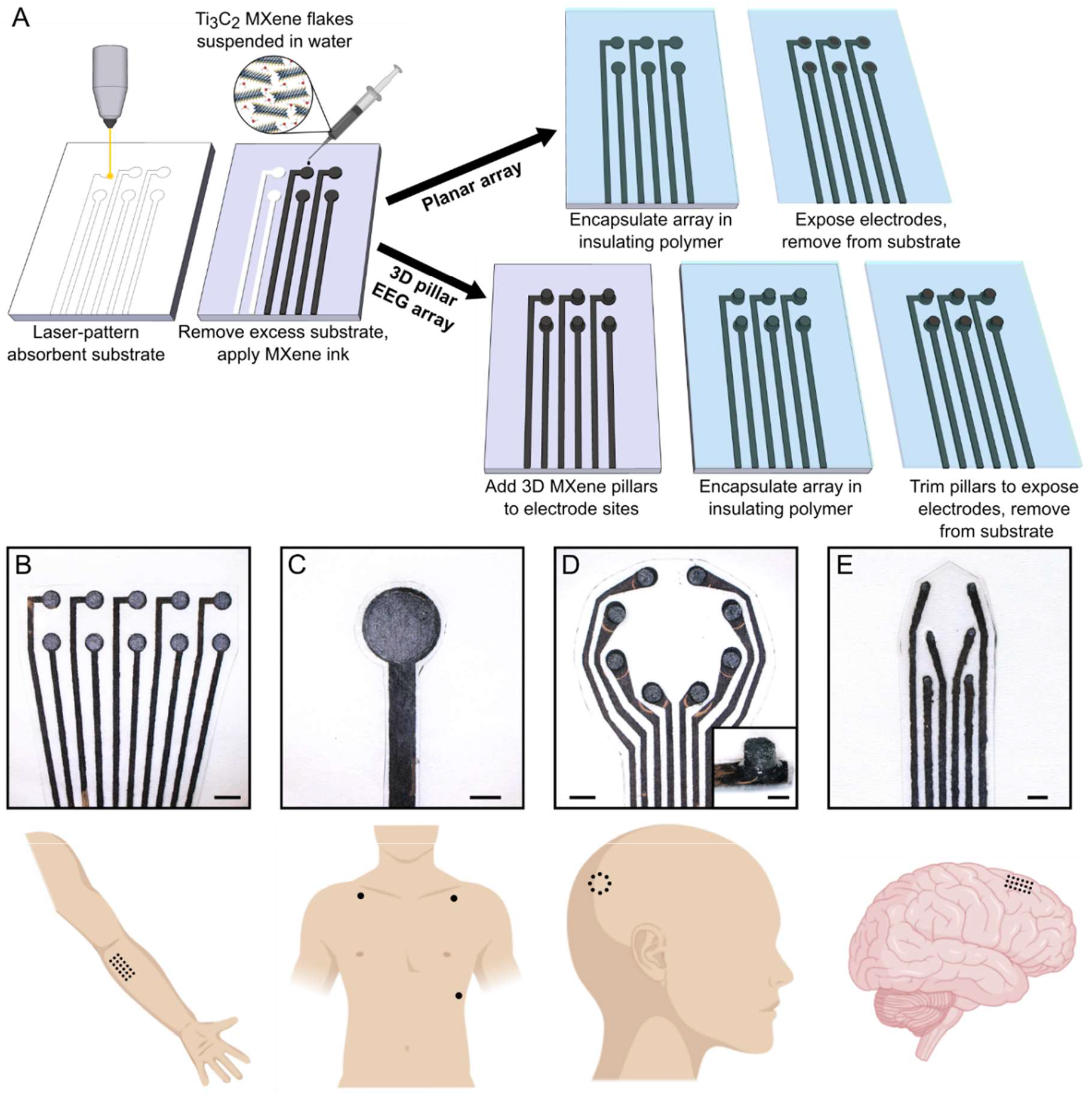
Rapid, high-throughput manufacturing of MXene ink-infused bioelectronics: MXtrodes. (**A**) Schematic of the fabrication method for laser-patterned planar and 3D mini-pillar MXene electrode arrays. (**B** to **E**) Photographs of different electrode array geometries (top) with their intended bioelectronic applications (bottom): (B) EMG, (C) ECG, (D) EEG, and (E) ECoG monitoring. Scale bars: (B to D) 5 mm; inset in (D) and (E) 2 mm.

### Electrical and electrochemical properties of MXtrodes

In the MXtrode arrays, the MXene-cellulose-polyester conductive composite forms the wires which carry the signal out to the recording amplifier. Thus, it is important that this composite is highly conductive in order to reduce Ohmic losses, minimize noise, and acquire high-quality signals. We measured the bulk conductivity of the MXene composites and determined it to be 3015 ± 333 S/m (fig. S3) which is in agreement with previously reported values for MXene-coated cellulose fibers (*50*). To highlight the conductivity advantage of Ti_3_C_2_ MXene compared to other conductive inks – which could in principle be used in our fabrication process – we also fabricated conductive composites using PEDOT:PSS and rGO inks with the same cellulose-polyester absorbent substrate. The bulk conductivity of PEDOT:PSS and rGO composites was 7.6 ± 0.4 S/m and 0.005 ± 0.002 S/m, respectively, both significantly lower than MXene.

To evaluate the impedance and charge transfer properties of MXene electrodes and compare them with other common bioelectronic materials, we conducted electrochemical measurements on MXtrodes with diameters ranging from 500 μm to 3 mm and compared them to 2.3 mm-diameter clinical Pt electrodes. Specifically, we performed electrochemical impedance spectroscopy (EIS), cyclic voltammetry (CV) and voltage transient (chronopotentiometry) experiments to measure the impedance magnitude, charge storage capacity (CSC), safe voltage window, and charge injection capacity (CIC) of each electrode, and to determine how these properties scale with electrode diameter. Data are shown in Table 1, and additional comparisons to other common electrode materials from literature are tabulated in table S1. EIS revealed that the MXtrodes of all diameters tested showed significantly reduced impedance compared to the Pt electrodes at frequencies below 500 Hz – where impedance is dominated by the electrochemical properties of the electrode interface (*51*) and where most physiologic signals of interest lie (Fig. 2A). At 10 Hz, the impedance of the MXtrodes ranging from 500 μm to 3 mm was 1343.3 ± 81.6 Ω (500 μm), 644.2 ± 97.3 Ω (1 mm), 451.2 ± 35.4 Ω (2 mm), and 241.4 ± 14.7 Ω (3 mm), while the impedance of the Pt electrodes was 8838.3 ± 1154.2 Ω (2.3 mm). At the reference frequency of 1 kHz – where the interface impedance is controlled by the solution resistance and electrode size (*51*) – the impedance of similar-sized MXtrodes is comparable to, and in some cases lower than, Pt and other bioelectronic materials like PEDOT:PSS (Fig. 2A and table S1). The 1 kHz impedance values can be found in Table 1. We attribute the exceptionally low impedance of the MXtrodes to the highly porous and rough morphology of the electrodes, which endows the interface with a high effective surface area.

**Fig. 2.**
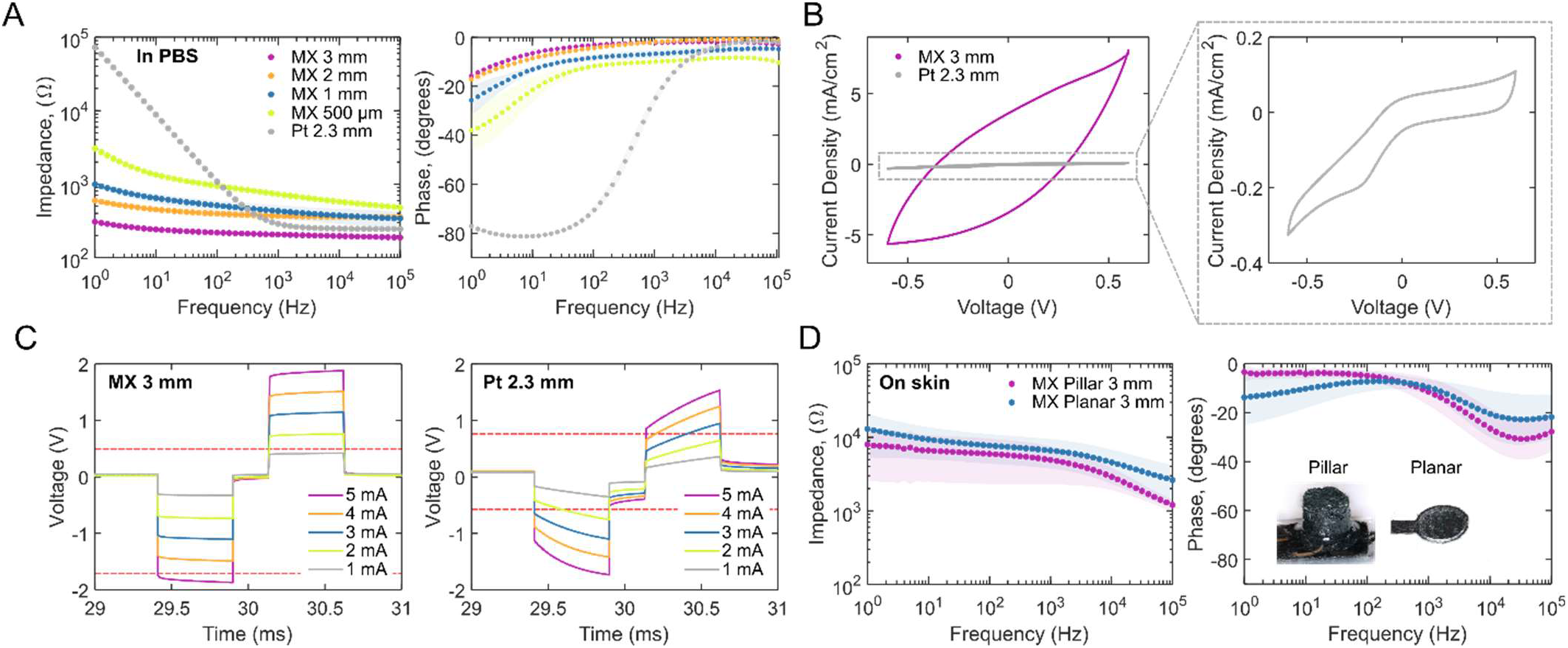
Electrochemical properties of MXtrodes. (**A**) EIS spectra measured in 1X PBS for 3 mm, 2 mm, 1 mm, and 500 μm planar MXene electrodes compared to 2.3 mm Pt electrodes. (**B**) CVs for 3 mm planar MXtrode and 2.3 mm Pt electrodes scanned from −0.6 – +0.6 V at 50 mV/s. (**C**) Voltage transients in response to biphasic current pulses, with *t_c_* = *t_a_* = 500 μs and *t_ip_* = 250 μs, current amplitudes ranging from 1 to 5 mA for 3 mm planar MXtrode and 2.3 mm Pt electrodes. Anodic and cathodic voltage limits for MXene and Pt are displayed on their respective plots as dashed red lines. (**D**) EIS spectra measured on skin for 3 mm MXtrode 3D pillar and planar electrodes.

**Table 1.** Summary of electrochemical properties of planar MXtrodes of varying diameters, compared to a 2.3 mm-diameter Pt electrode contact.

| Material | Electrode diameter | 1 kHz $\vert Z\vert $ in PBS ($\Omega$) | Potential Limits vs. Ag/AgCl (V) | CSC<sub>c</sub> (mC cm<sup>-2</sup>) | CIC<sub>c</sub> (mC cm<sup>-2</sup>) |
| --- | --- | --- | --- | --- | --- |
| MXene | 3 mm | 205.6 ± 11.1 | -1.8 – 0.6 | 233.1 ± 19.6 | 0.55 ± 0.10 |
|  | 2 mm | 369.2 ± 40.2 | -1.8 – 0.6 | 392.3 ± 9.8 | 0.69 ± 0.03 |
|  | 1 mm | 430.2 ± 86.1 | -1.8 – 0.6 | 604.8 ± 61.4 | 1.07 ± 0.07 |
| | 500 μm | 729.2 ± 71.2 | -1.8 – 0.6 | 1024.1 ± 3.0 | 2.12 ± 0.48 |
| Pt | 2.3 mm | 287.3 ± 7.8 | -0.6 – 0.8 | 4.3 ± 0.2 | 0.07 ± 0.003 |

The safe voltage window for MXtrodes determined from wide-scan CVs is −1.8 – +0.6 V (fig. S4, A and B), showing that MXene is exceptionally stable in the cathodic region, with hydrolysis beginning at −1.9 V. This wide safe potential range is advantageous for therapeutic electrical stimulation applications, and while the safe anodic potential limit (+0.6 V) is slightly lower than that of Pt, PtIr, or IrOx (+0.8 V), stimulation waveforms can be engineered to minimize voltage excursions in the anodic range while taking advantage of the large cathodic limit (*52–54*). While previous studies have reported anodic oxidation of Ti_3_C_2_ films at +430 mV (*55*), we observed no oxidation peak and negligible current loss when MXtrodes were cycled within the range of −1.8 – +0.6 V. We attribute the improved anodic stability to the rough surface morphology of the MXtrodes, resulting in decreased current density. In the cathodic region, the 1 mm, 2 mm, and 3 mm MXtrodes showed stable capacitive behavior down to −1.8 V, however the 500 μm-diameter MXtrode showed initial evidence of faradaic current near this negative limit (fig. S4C) likely due to size effects. Still, the 500 μm-diameter MXtrode showed stable behavior through repeated CV cycles and no evidence of hydrolysis, confirming −1.8 V as a reasonable cathodic limit.

Analysis of cathodal CSC (CSC_C_) for MXtrodes and Pt electrodes from CVs at 50 mV/s within their respective water windows reveals enhanced capacitive charging and charge delivery properties for MXtrodes with ~100 times larger CSC_C_ than Pt electrodes (Table 1) and other common materials such as PtIr and PEDOT:PSS (table S1). To enable a more direct comparison of CSC_C_ values between MXtrodes and Pt electrodes, we also ran CV scans in the intersection of the MXene and Pt voltage windows, −0.6 – +0.6 V (Fig. 2B and fig. S4D). Even in this more constrained voltage range, the MXtrodes showed more than 20 times enhanced CSC_C_ compared to Pt, which we attribute both to the exceptionally high intrinsic capacitance of Ti_3_C_2_ MXene (*56–58*) and to the high effective surface area of the MXtrode surface (table S2). The scaling dependency of CSC_C_ on the electrode diameter for the MXtrodes is shown in fig. S4E. The non-linear relationship between CSC_C_ and electrode diameter is expected, and reflects the known phenomenon of electrochemical charge exchange happening predominantly at the edge of the electrode (*54, 59*).

Finally, we measured the voltage transients evolved on each electrode when used to deliver charge-balanced, cathodal-first biphasic current pulses ranging from 1 to 5 mA with a duration of 500 μs in both phases (*t_c_, t_a_*) and an interpulse interval (*t_ip_*) of 250 μs (Fig. 2C and fig. S4, F to I). For the 500 μm-diameter MXtrode, the current amplitude range was restricted to 600 μA – 2 mA. The maximum cathodal excursion potential, *E_mc_*, was taken 10 μs after the cathodal pulse end and CICC was defined as the injected charge at which *E_mc_* would reach the water reduction potential. The resulting CICC values, shown in Table 1, reveal that the MXtrodes significantly outperform Pt electrodes, with MXtrodes showing ~10 times larger CIC_C_ than the Pt electrodes. Furthermore, we found a similar ~10 times improvement in CIC_C_ when we compared MXtrodes to other common electrode materials (table S1). This result has significant implications for stimulation applications, and suggests that MXtrodes may offer more efficient charge transfer than current state-of-the-art Pt electrodes. This could potentially prolong battery life for implantable simulation systems such as deep brain stimulation (DBS), vagal nerve stimulation (VNS), and cardiac pacemakers. The scaling dependency of CICC on electrode diameter for the MXtrodes is shown in fig. S4J, again revealing the expected non-linear scaling dependency resulting from edge effects. A schematic depicting the relative sizes of the MXtrodes included in the analysis is shown in fig. S4K.

While measurements in saline allow comparing the properties of MXtrodes to current standard electrode materials such as Pt, for epidermal sensing applications it is also essential to evaluate the impedance at the interface with human skin. Specifically, achieving low electrode-skin impedance is key for recording high-fidelity signals (*60, 61*) and becomes particularly challenging in gel-free structures like MXtrodes. Thus, we measured EIS for 3 mm-diameter MXtrodes, both in planar and 3D configurations, on clean human skin following standard preparation with an alcohol swab and light abrasion with 3M Trace Prep tape. At 1 kHz, the planar and 3D MXtrodes showed impedances of 6.6 ± 2.9 kΩ and 4.9 ± 2.6 kΩ, respectively, with the lower impedance of the 3D electrodes attributable to the improved contact from the protruding mini-pillars pressing into the skin (Fig. 2D). When normalized by their geometric surface area (GSA) of 0.071 cm^2^, the impedance is 0.47 ± 0.20 kΩ·cm^2^ for planar and 0.35 ± 0.12 kΩ·cm^2^ for 3D MXtrodes, which are among the best values reported so far for dry, gel-free epidermal electrodes (*1, 62, 63*) and ~100 times lower than those of commercially available gelled Ag/AgCl electrodes commonly used for electrodiagnostics and monitoring (*45*).

### Epidermal sensing of bioelectric signals in humans

Motivated by the exceptionally low electrode-skin interface impedance of the dry MXtrodes, we investigated their use in a variety of human epidermal sensing applications, with custom geometries specifically designed for each application. First, we acquired scalp EEG on a healthy human subject using high resolution gel-free MXtrodes and standard gelled Ag/AgCl EEG electrodes for comparison. We designed an 8-channel MXtrode, with 3 mm-diameter 3D mini-pillar electrodes arranged in a ring around a central opening, where we placed a standard 1 cm-diameter gelled Ag/AgCl EEG electrode for side-by-side comparison of simultaneously acquired EEG (Fig. 3A). In the first EEG task, we placed the MXtrode array over the parietal region near EEG site P1 (as defined in the EEG 10-20 system, located over inferior parietal cortex to the immediate left of the midline). The gelled Ag/AgCl EEG electrode was placed in the center of the MXtrode ring (Fig. 3B). Ground and reference for all EEG recordings were gelled Ag/AgCl electrodes placed at the center forehead and left mastoid, respectively. The slightly lateralized recording position was chosen because the subject had the shortest hair at that location (~5 mm). Before placing the electrodes on the skin, the entire recording area was cleaned with an alcohol swab and lightly abraded with 3M Trace Prep tape. Notably, we found that the electrode-skin interface impedance at 1 kHz for the dry MXtrodes was 2.8 ± 0.9 kΩ, while the impedance of the larger gelled Ag/AgCl electrode during the same experiment was 1.2 kΩ at 1 kHz (Fig. 3C). Given the critical role of the electrode-skin interface impedance in determining the quality of scalp EEG signals (*64*), most standard EEG electrodes require conductive gels at this interface, as well as a large contact area of at least ~1 cm^2^ to achieve suitably low impedance (e.g., less than 5 kΩ) (*65*). Due to their enhanced material and surface area properties, our mm-scale, gel-free MXtrodes can achieve strikingly low impedance, which enables high-resolution EEG recording. We recorded EEG in 2 min sessions, with the subject alternating between a resting state with the eyes closed and a resting state with the eyes open and fixating on a target. In both tasks, the EEG signal recorded on the dry MXtrodes was indistinguishable from the signal recorded on the gelled Ag/AgCl electrode (Fig. 3D). Additionally, a clear 10 Hz alpha rhythm emerged in the eyes closed state with a significantly higher amplitude than in the eyes open condition (Fig. 3, E and F). This alpha signature is one of the most reliable and widely studied behaviorally-linked EEG signatures in human subjects research (*66*), and arises from endogenous thalamic input to the visual cortex in the absence of visual input (i.e. when the eyes are closed) (*67*). During the recording session, there was no significant difference between the alpha bandpower on the gelled Ag/AgCl electrode and any individual dry MXtrode, confirming that the signals were comparable between the electrode types (fig. S5). Interestingly, when alpha bandpower values were calculated in 1 s windows with 0.5 s of overlap and observed sequentially, distinct spatiotemporal patterns of alpha activation emerged even across the small sampled scalp area (movie S1). MXtrodes could be spaced much closer than they were in the ring configuration we used in this experiment, however the density we tested already approximates or exceeds the densest configurations of traditional Ag/AgCl electrodes in common use today (the 10-5 system). This highlights the potential of mm-scale, gel-free MXtrodes to enable ultra high-density EEG mapping.

**Fig. 3.**
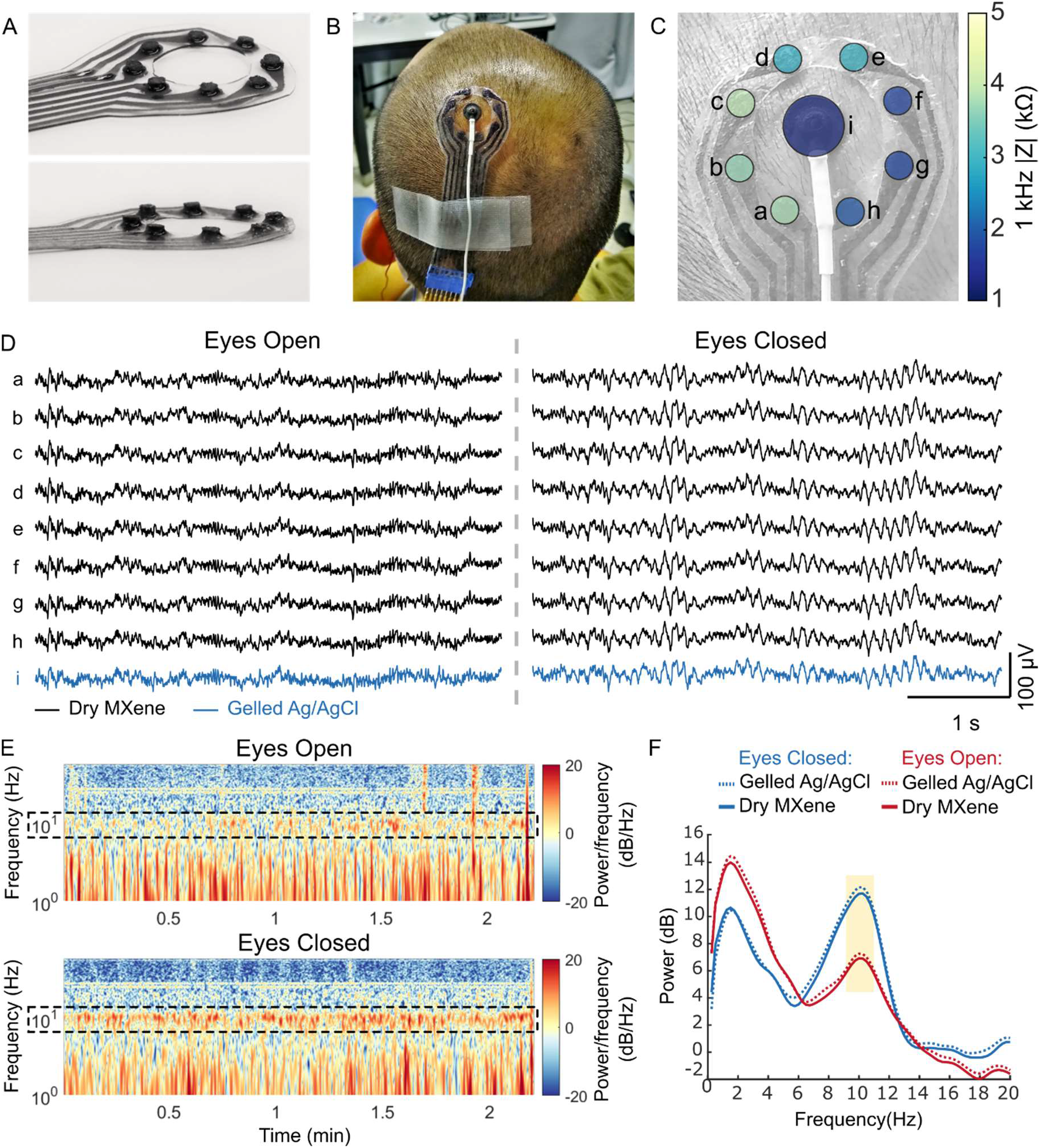
Dry EEG recording enabled by 3D pillar MXtrodes. (**A**) Images of a MXtrode 3D EEG array with eight 3 mm-diameter MXene electrodes in a circular arrangement around a central opening. (**B**) Image of MXtrode electrode array and standard gelled Ag/AgCl cup electrode placed on head of human subject. (**C**) Map of 1 kHz impedance values for all electrodes on the subject’s head. (**D**) Segments of recorded EEG signal from all electrodes during the eyes open (left) and eyes closed (right) tasks at resting state. (**E**) Spectrograms of the EEG signal recorded on MXene electrode b in the eyes open (top) and eyes closed (bottom) conditions. Alpha frequency band is enclosed in dashed box to highlight differences between eyes open and eyes closed states. (**F**) Power spectral density during eyes open and eyes closed EEG recordings. The 8–12 Hz alpha band is highlighted.

For the second EEG task, the electrodes were removed and replaced at site C3 near the hand motor area. The precise location of the hand motor area was determined using single pulses of transcranial magnetic stimulation (TMS) to evoke lateral finger movements, and the electrodes were centered over this location (fig. S6A). The subject performed 2 min periods of imagined and actual hand flexion while the EEG was recorded simultaneously with the co-located MXtrodes and gelled Ag/AgCl electrode. During this motor task, we also found that the EEG signal was indistinguishable between the two electrode types. Furthermore, we observed a suppression of the 8-12 Hz motor mu rhythm during the actual hand flexion relative to the imagined hand flexion (fig. S6B). The mu suppression signature during imagined hand movement is an important EEG feature which has been successfully used as a control signal for EEG-based braincomputer interfaces (BCIs) (*68, 69*). Together, the results of these EEG experiments confirm that mm-scale, gel-free MXtrodes can record EEG signal at least as well as standard gelled Ag/AgCl EEG electrodes, while also improving spatiotemporal resolution for high-density EEG applications. Furthermore, the application of MXtrodes to the scalp is faster and easier than standard Ag/AgCl electrodes and leaves no residue on the head. This positions MXtrodes to dramatically lower the effort of conducting EEG experiments, and to expand the use of high-fidelity EEG to environments outside the laboratory.

Next, we evaluated MXtrodes for high-density surface electromyography (HDsEMG) recording, muscle activation mapping and localization of the of innervation zones (IZs). HDsEMG is attracting growing interest for a number of applications in neuromuscular diagnostics and rehabilitation, including control of multifunctional prostheses (*19*), studies of muscle activation and coordination (*70*), peripheral nerve/muscle fiber conduction velocity measurements (*71*), and for accurate localization of neuromuscular junctions (NMJs) to target chemodenervation therapies for muscle spasticity (*72, 73*). HDsEMG recordings require flexible, large-area, and high-density electrode arrays capable of covering the wide range of muscle sizes. To demonstrate the versatility of the MXtrode fabrication process, we created custom HDsEMG arrays to map muscle activation and localize IZs in two muscle groups of different sizes (Fig. 4). First, we used a 20-ch planar MXtrode array placed over the *abductor pollicis brevis* (APB) at the base of the thumb. In this experiment, the 3 mm-diameter dry MXtrodes had an average electrode-skin interface impedance of 54.6 ± 28.4 kΩ at 1 kHz (fig. S7A). We then stimulated the median nerve with a handheld bipolar stimulator to evoke APB contractions and recorded the EMG on the MXtrode array. We calculated the mean evoked muscle response across stimulation trials (Fig. 4A) and constructed a latency map of the peak of the evoked response. The location with the shortest latency indicates the location of the IZ, which can be seen overlaid on the subject’s hand in Fig. 4B, and it is in good agreement with expectations based on anatomical landmarks (*74*).

**Fig. 4.**
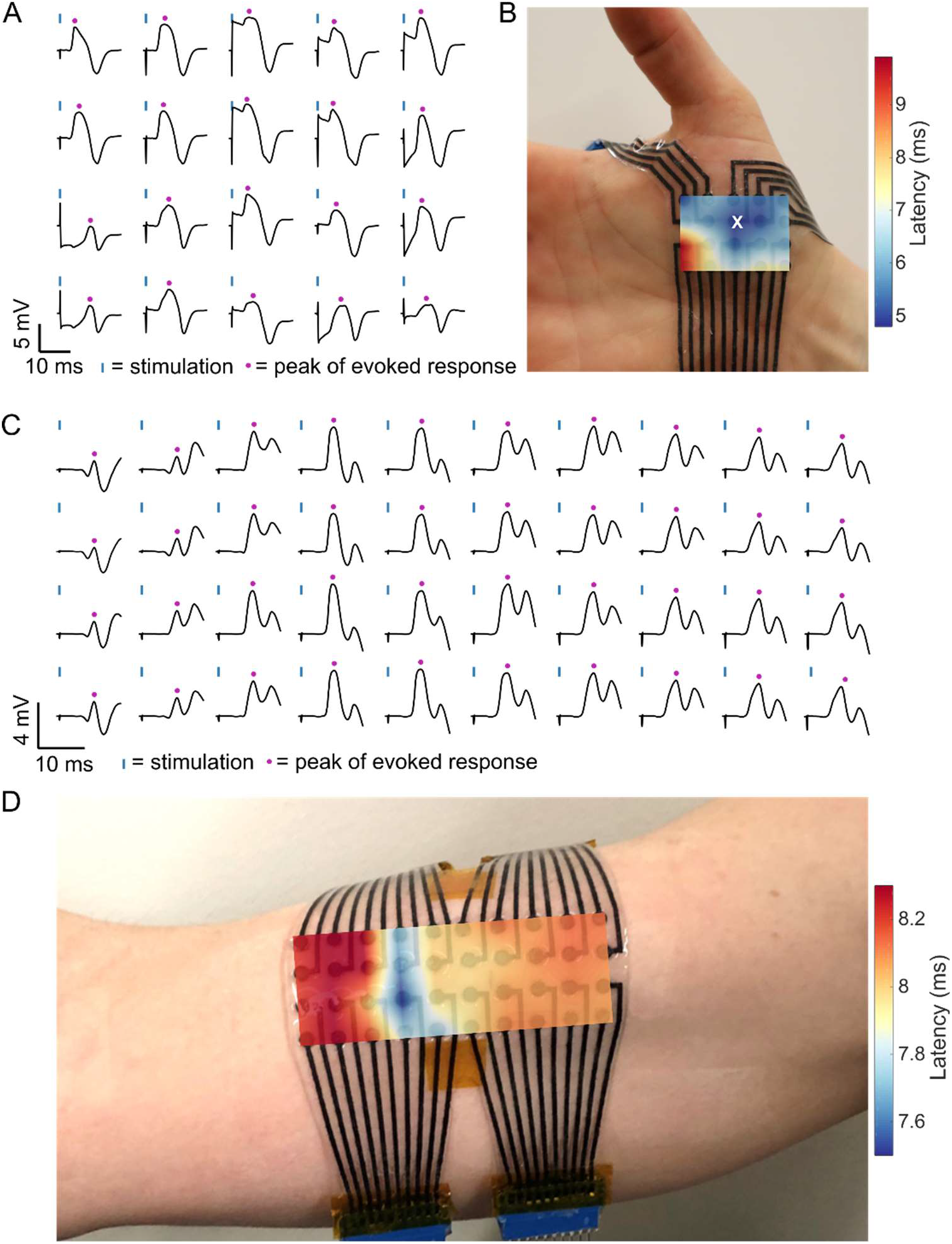
High density surface EMG mapping with MXtrode arrays. (**A** and **B**) EMG recordings from the APB muscle. (A) Average evoked response following N=10 median nerve stimulation epochs recorded on 20-ch MXtrode array placed over APB. Blue ticks indicate time of nerve stimulation, and purple dots indicate time of peak evoked response. (B) Latency map of peak response overlaid on photo of the MXtrode array on the APB. White “x” indicates the channel with shortest latency, corresponding to the IZ. (**C** and **D**) EMG recordings from the biceps muscle. (C) Average evoked response following N=10 supraclavicular nerve stimulation epochs recorded on 40-ch MXtrode array placed over the biceps. Blue ticks indicate time of nerve stimulation, and purple dots indicate time of peak evoked response. (D) Latency map of peak response overlaid on photo of the MXtrode array on the subject’s biceps. Distributed IZ running perpendicular to the muscle is apparent as the band with the shortest latency.

Following the APB mapping experiment, we mapped the activation of the larger *biceps brachii* with a 40-ch planar MXtrode array. Here, the 3 mm-diameter dry MXtrodes had an average electrode-skin interface impedance of 22.0 ± 14.3 kΩ at 1 kHz (fig. S7B). Unlike the smaller APB muscle, which has a small and spatially confined IZ corresponding to a single NMJ, the larger *biceps brachii* has distributed NMJs in IZ regions that run perpendicular to the muscle and are typically located near the center (*75*). To further confirm the validity of our approach, we used two different methods to localize these IZs in the biceps: first we stimulated the supraclavicular nerve and constructed a latency map of the peak evoked response, similar to the methods described for the APB (Fig. 4, C and D). This produced a clear mapping of the IZ location as the region with the shortest latency, running across the short head of the *biceps brachii*. Second, we recorded motor unit action potentials (MUAPs) as the subject performed isometric contractions of the biceps (fig. S7, C and D). In these recordings, bipolar subtraction of the raw EMG signal along the length of the biceps muscle revealed MUAPs that propagated outward in both directions from the IZ, with signal inversion and a clear latency as the MUAP traveled away from the IZ (fig. S7D). Localization of the biceps IZ obtained from both methods were in excellent agreement with each other, and with previous reports (*75*). These results demonstrate that dry, high-resolution MXtrode arrays are capable of mapping muscle activation with high accuracy in order to precisely localize IZs in both small and large muscle groups.

To demonstrate the applicability of MXtrodes to electrocardiogram (ECG), we acquired ECG recordings on a healthy human subject with 1.3 cm-diameter MXtrodes in a simplified 3-electrode montage as shown in Fig. 5A. For validation and signal comparison, we recorded sequentially from the MXtrodes and from 1 cm-diameter pre-gelled Ag/AgCl electrodes, placed in the same locations. The dry MXtrodes and pre-gelled electrodes had average 1 kHz skin-electrode impedances of 1.29 kΩ and 1.38 kΩ, respectively. On both types of electrodes, characteristic ECG features were clearly visible, with the P wave followed by the QRS complex and the T wave (Fig. 5, B and C). Mean R peak amplitude, however, was slightly higher on the MXtrodes compared to the commercial electrodes: 2.55 ± 0.06 mV vs. 2.47 ± 0.30 mV. Thus, we confirmed that dry MXtrodes can record the ECG with comparable signal quality compared to standard clinical gelled electrodes.

**Fig. 5.**
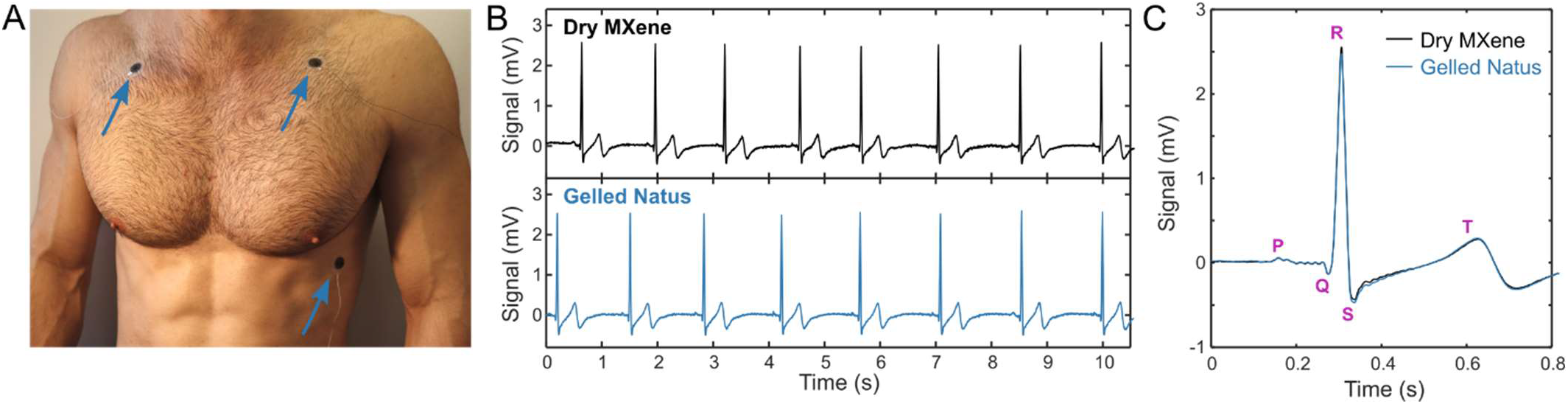
Electrocardiography with MXtrodes. (**A**) Photo of ECG recording setup on human subject. Electrodes were interchanged in the same locations to obtain sequential recordings from either dry MXene or pre-gelled commercial electrodes. (**B**) Ten seconds of ECG recordings on the dry MXtrodes (top) and the pre-gelled commercial electrodes (bottom). (**C**) Average ECG waveforms recorded on the two electrode types, marked with salient ECG features.

As a final demonstration of MXtrodes for human epidermal sensing, we acquired electrooculography (EOG), which has applications in ophthalmological diagnosis, BCIs, and for monitoring attention and fatigue (*76, 77*). The EOG signal arises from the standing dipole potential between the positively charged cornea and the negatively charged retina, which enables tracking eye movements as this dipole is rotated. With the same 1.3 cm-diameter dry MXtrode geometry used in ECGs, we recorded the EOG in two configurations to track up-down and left-right eye movements. By placing MXtrodes above and below the eye, recorded voltage fluctuations could be decoded to track the up and down movements of the eye (fig. S8, A and B). Similarly, placing MXtrodes on both sides of the eyes enabled decoding the left-right movements (fig. S8, C and D).

### Cortical neural recording and microstimulation with MXtrodes

Beyond epidermal sensing modalities, the favorable electrochemical interface of the MXtrodes also supports their use for implantable sensing and stimulation applications. One such application is intraoperative electrocorticography (ECoG), a common mapping technique used in resective brain surgery for epilepsy or tumors. We acquired ECoG recordings in an anesthetized swine, a relevant model system in neuroscience given its gyrencephalic structure and neuro-anatomical similarity to the human brain. In this experiment we inserted a 6-ch array of 500 μm-diameter planar MXtrodes through an 8 mm craniotomy/durotomy and placed MXtrodes in direct contact with the cortical surface (Fig. 6A). The array configuration consisted of 3 rows of electrode pairs with 5 mm inter-row spacing and 4.5 mm pitch, so that the rows of electrodes spanned several cortical gyri. A few seconds of representative raw ECoG signal are shown in Fig. 6B. The signals were large in amplitude with negligible 60 Hz noise interference, as evidenced by the power spectra (Fig. 6C). Furthermore, maps of interpolated voltage across the MXtrode array revealed stereotyped spatial patterns emerging during the “up” and “down” states in the ECoG signal (Fig. 6, D and E), highlighting the advantages and opportunities offered by high-density cortical brain mapping with MXtrodes.

**Fig. 6.**
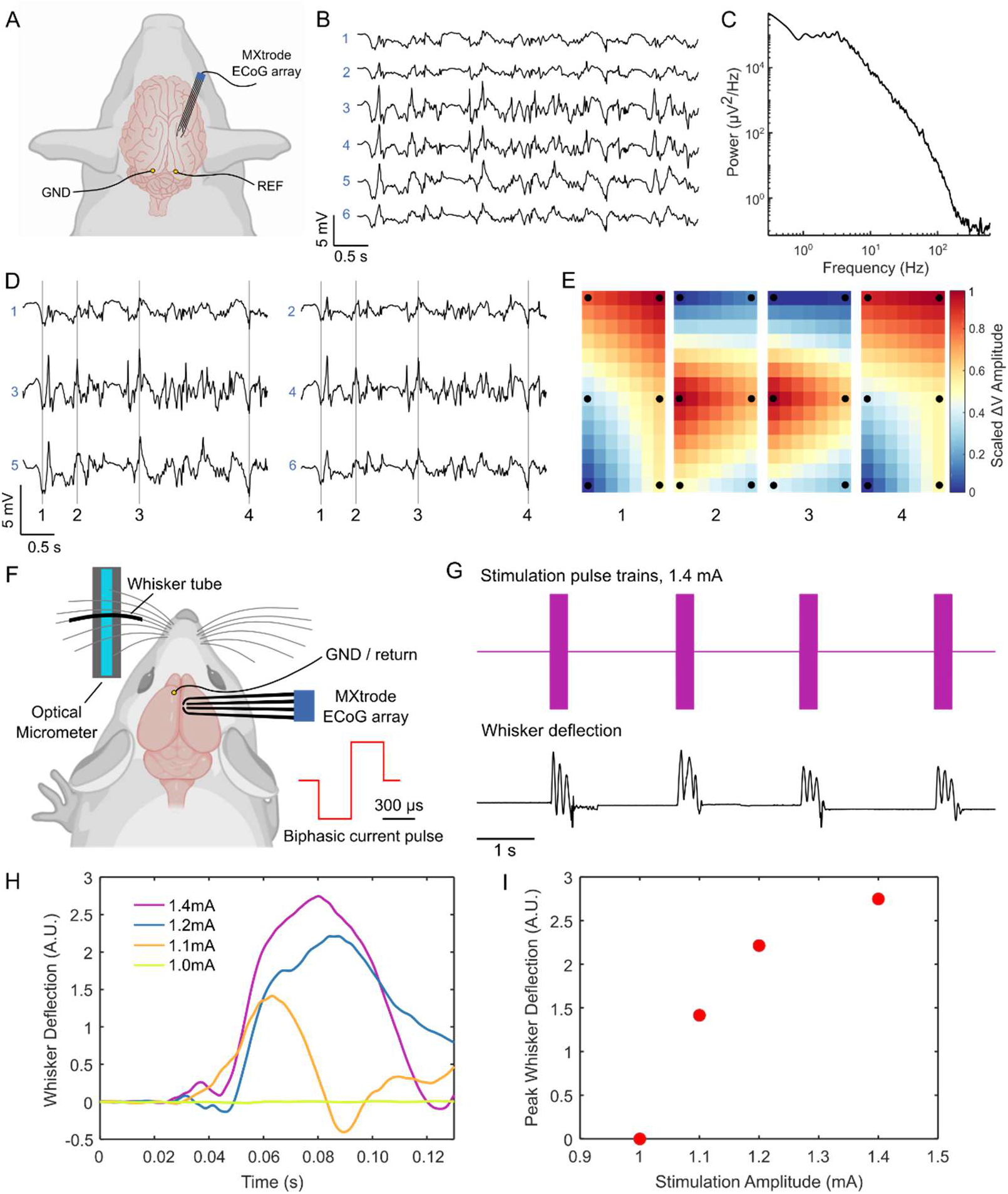
ECoG recording and cortical stimulation with MXtrode arrays. (**A**) Schematic depicting ECoG recording setup with the 6-ch array of 500 μm-diameter MXtrodes placed subdurally on somatosensory cortex. (**B**) A few seconds of representative ECoG data recorded on the MXtrode array. (**C**) Power spectral density of the ECoG recording, illustrating the low-noise quality of the ECoG signals obtained, evidenced by the lack of a 60 Hz noise peak. (**D**) Segment of ECoG data, displayed according to the spatial arrangement of the 6 MXtrodes. (**E**) Instantaneous snapshots of the voltage mapping recorded across the 6 MXtrodes reveals stereotyped patterns during down states (panels 1 and 4) and up states (panels 2 and 3). The timing of these voltage snapshots is indicated in (E) by the vertical lines. Voltage was interpolated across the array and normalized, with black dots indicating location of the 6 MXtrode contacts. (**F**) Schematic of the cortical stimulation setup, with the 4-ch array of 500 μm-diameter MXtrodes placed over barrel cortex, and the optical micrometer used to detect and amplify the whisker deflection signal. (**G**) Whisker deflection data recorded by the optical micrometer during a series of stimulation pulse trains delivered at 1.4 mA. (**H**) Average first whisker deflection for each stimulation pulse amplitude, time-aligned by the stimulation onset. (**I**) Whisker deflection amplitude scales with stimulation amplitude, with stimulation at 1.0 mA falling below the threshold required to evoke whisker movement.

In addition to ECoG recording, direct stimulation of the cortical surface is used clinically for intraoperative cortical mapping (*78*) and neuromodulation therapies (*79*), as well as for closed-loop BCIs (*80*). Given that the MXtrodes showed superior CSC and CIC compared to Pt, a material commonly used in stimulating electrodes, we sought to demonstrate the effectiveness of MXtrodes for electrical stimulation by evoking motor responses via intraoperative stimulation in rat brain. Specifically, we placed a 500 μm-diameter planar MXtrode subdurally onto the whisker motor cortex of an anesthetized rat. Contralateral to the MXtrode, an optical micrometer was positioned to track whisker displacement, with one whisker placed into a plastic tube to maximize detection sensitivity (Fig. 6F). Trains of charge-balanced, cathodal-first stimulation pulses with amplitudes ranging from 1.0 to 1.4 mA were delivered through the MXtrode. We observed stimulation-evoked whisker movements for amplitudes greater than 1.0 mA, with the whisker deflection amplitude scaling with the stimulation intensity (Fig. 6, G to I and movie S2). Whisker movements registered as oscillatory deflections on the micrometer, with the first deflection peak always being the largest in amplitude. To compare whisker deflection across stimulation current amplitudes, we computed the mean amplitude of the first whisker deflection peak for each stimulation trial at each current level. These results confirm that MXtrodes are capable of delivering electrical stimulation to effectively modulate neural activity.

### Compatibility of MXtrodes with clinical imaging

With the widespread adoption of bioelectronic technologies, compatibility with clinical imaging has become increasingly important. MRI and CT are the two most common imaging techniques used in the diagnosis of injury and disease, as well as in image-guided interventions. However, many of the conductive materials traditionally used in bioelectronic devices are incompatible with the challenging MRI environment and can produce heating or exert forces on the tissues. Even devices considered MR-safe often produce imaging artifacts that shadow the surrounding anatomical structures because of the mismatch in magnetic susceptibility between the device materials and the tissue (*81*). These challenges are compounded at high field strengths, which are increasingly being used for high resolution imaging, as well as for novel functional and metabolic imaging techniques (*82, 83*). While the magnetic susceptibility of Ti_3_C_2_ MXene had not been previously characterized, we hypothesized from the weak dia- and paramagnetic properties of C and Ti, respectively, that Ti_3_C_2_ may have a low magnetic susceptibility and thus prove compatible with the MR environment. To verify our hypotheses, we performed MRI scans of MXtrodes and measured the magnetic properties of Ti_3_C_2_ at body temperature. First, we imaged cross-sections of 3 mm-diameter planar MXtrodes and 2.3 mm-diameter commercial Pt electrodes embedded in conductive agarose phantoms in a 9.4T high-field research MRI system (Fig. 7A). In the MR images, we found significant shadowing around the Pt electrodes, while no artifact was visible around the MXtrode (Fig. 7B). In fact, the MXene composite forming the conductive electrode was almost completely indistinguishable from the surrounding PDMS encapsulation. To further explore the MRI compatibility of the MXtrodes, we next imaged an array of 3 mm-diameter 3D mini-pillar MXtrodes – the same array used for EEG recording in this work – in a 3T clinical MRI scanner. The MXtrodes were placed atop a tissue phantom and imaged using a battery of scan sequences typical of structural and functional MRI. Regardless of the scan sequence, the MXtrodes showed no artifact and were nearly invisible in the images (fig. S9A). Thermal IR images of the MXtrode array immediately following a 10 min scan sequence also showed no evidence of RF heating (fig. S9B). Finally, we measured the magnetic susceptibility of a Ti_3_C_2_ free-standing film at 310 K, and found it to be 2.08 x 10^-7^ (fig. S9C), indicating that Ti_3_C_2_ is weakly paramagnetic with magnetic susceptibility very close to that of human tissues (−11.0 to −7.0 x 10^-6^). This innate property of Ti_3_C_2_, which has not been reported previously, leads directly to the excellent compatibility of the MXtrodes with MRI imaging. For comparison, the magnetic susceptibility of Pt is 2.79 x 10^-4^, several orders of magnitude larger than the susceptibility of human tissues(*81*). In situations where it might be important for implanted electrodes to be visible in MRI images, such as in epilepsy seizure foci localization relative to electrode locations, a marker such as iron oxide nanoparticles could in principle be incorporated into the electrode contacts to aid in their visualization (*84*).

**Fig. 7.**
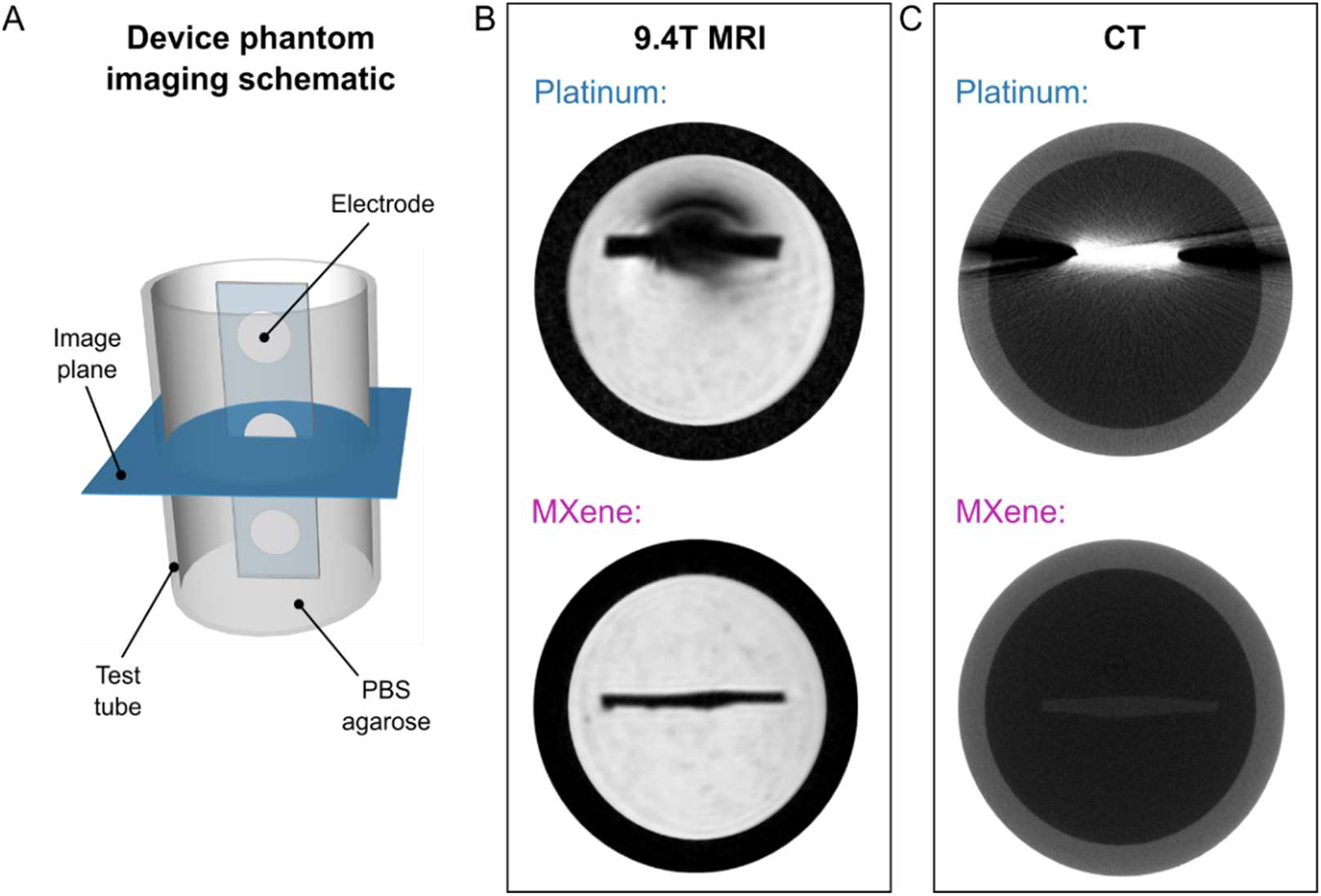
MRI and CT compatibility of MXtrodes. (**A**) Schematic of the phantom imaging setup for MRI and CT imaging. Strips of disk electrodes (3 mm-diameter MXtrodes or 2.3 mm-diameter clinical Pt ECoG strip electrodes) were embedded in a conductive agarose phantom and imaged in the transverse plane. (**B**) A high-field 9.4T MRI scan showing significant shadow artifact and image distortion around the Pt electrode (top), but no visible artifact from MXtrodes (bottom). The MXene contact is indistinguishable from the PDMS encapsulation in the device. (**C**) High-resolution CT scans with Pt electrodes (top) showing significant x-ray scattering artifacts, while no artifact is visible from the MXtrodes (bottom).

As for CT compatibility, the X-ray attenuation characteristics resulting from the high density and large atomic mass of metals in conventional electrodes poses challenges for CT imaging with bioelectronic devices (*85*). Multilayer Ti_3_C_2_ MXene has a density of about 4 g/cm^3^, which is ~5 times lower than Pt. Thus, we hypothesized that Ti_3_C_2_ could minimize attenuation and scattering artifacts in CT (*86*). To test our hypothesis, we imaged 3 mm-diameter planar MXtrodes and 2.3 mm-diameter commercial Pt electrodes embedded in conductive agarose phantoms in a microCT scanner. As expected from the density matching considerations, we observed significant X-ray scattering artifacts around the Pt electrodes, but not around the MXtrodes (Fig. 7C).

## Discussion

In this work we developed a new class of MXene-based bioelectronic interfaces and demonstrated their use for high-fidelity recording and effective stimulation of neural and neuromuscular circuits across multiple scales. The simple fabrication method reported here offers a scalable and low-cost means of producing large-area, multichannel electrode arrays from high performance, solution processable 2D materials. The method is conducive to large-scale manufacturing, a key aspect for translation beyond the lab and into clinical and consumer markets. It also enables rapid customization of array geometries for various applications as well as for patient- or subject-specific fit where desired. The exceptional properties of Ti_3_C_2_ endow MXtrodes with impedance and charge delivery properties that meet and even exceed current state-of-the-art noble metal electrode materials in both epidermal and implantable use cases. In epidermal sensing applications, the remarkably low electrode-skin interface impedance of gel-free MXtrodes opens exciting new possibilities for high-resolution EMG and EEG, while eliminating the challenges associated with wet conductive gels. In HDsEMG mapping, MXtrode arrays hold promise for accurately localizing NMJs for use with neurolytic procedures for the treatment of spasticity and excessive muscle tone. In the future, this may eliminate the need for intramuscular needle EMG stimulation in search of the NMJ – the optimal place for injecting medications (e.g. botulinum toxin and phenol) for denervation. Furthermore, such HDsEMG arrays can also be integrated within an active prosthesis system where EMG from the residual limb serves as input signal for the prosthesis control (*87, 88*). In EEG applications, a dry electrode system enabled by MXtrodes may offer a route toward minimizing skin breakdown and alleviating many of the key logistical challenges associated with current gelled EEG systems, such as the time required to apply each electrode and impedance fluctuations over time as the gels dry out. These advantages, coupled with the possibility for obtaining high density EEG recording with mm-scaled MXtrodes, also make this technology attractive for neuroscience research (*89, 90*), non-invasive BCI systems (*91, 92*), neuro-rehabilitation (*93*), and brain-controlled videogaming (*94*). The compatibility of MXtrodes with both CT and MRI imaging enables accurate, artifact-free imaging without the need for removing electrodes from the subject before the scan, thus opening exciting possibilities for multimodal studies in both clinical diagnostics and neuroscience research, such as those combining simultaneous EEG with functional MRI or GluCEST imaging (*83, 95*). In addition to the low interface impedance, the extensive characterization of the electrochemical behavior of MXtrodes presented in this work demonstrates for the first time the advantages of Ti_3_C_2_ for electrical stimulation, as highlighted by remarkably high CSC and CIC compared to many commonly adopted bioelectronic materials. This finding suggests that MXtrodes may represent a suitable alternative for stimulation electrodes with enhanced safety, charge transfer efficiency, and reduced power consumption. While the technology presented here already shows great potential for translation, ongoing efforts include evaluating and improving the stability of Ti_3_C_2_ for long-term use (*47, 96, 97*), optimizing the MXene infusion process to maximize the MXene loading and conductivity of the resulting composite (*50*), and exploring alternative encapsulation materials to further improve the flexibility and breathability of the electrode arrays (*19*). Overall, our results indicate that MXene-based bioelectronics have tremendous potential for enabling the next generation of soft interfaces for advanced healthcare diagnostics, monitoring and therapy, as well as for research and consumer electronics.

## Materials and Methods

### Synthesis of Ti_3_C_2_ MXene

Ti_3_C_2_ MXene was provided by Murata Manufacturing Co., Ltd. It was produced using the MILD synthesis method (*49*) to create an ink of 30 mg/mL Ti_3_C_2_ in deionized water, which was placed in a vial and sealed under Argon atmosphere for long term storage.

### Fabrication of MXtrode devices

Devices were fabricated by first laser-patterning absorbent nonwoven textile substrates comprised of hydroentangled 55% cellulose / 45% polyester blend (Texwipe TechniCloth) using a CO_2_ laser (Universal Laser Systems PLS 4.75) such that electrode array patterns were easily separable from the surrounding textile, but could still be lifted and handled as one sheet. These were transferred to a thin and slightly tacky bottom layer of 1:10 PDMS (Sylgard 184) on a flat acrylic sheet, and the excess textile surrounding the array patterns was then peeled up. The textile patterns were then inked with 30 mg/mL Ti_3_C_2_ MXene, allowed to air dry for 15 min, then placed in a vacuum oven (Across International) at 70 °C and 60 mmHg for 1 h to remove all remaining water. For devices incorporating 3D mini-pillar electrodes, 3 mm circles were cut from absorbent cellulose sponges (EyeTec Cellulose Eye Spears) using a 3 mm biopsy punch and these circular sponges were inked with MXene and placed onto the electrode locations of the MXene-textile constructs at the same time that the laser-patterned textile was inked. The subsequent drying steps were identical. No adhesive was necessary at this interface, as the dried MXene and the subsequent PDMS encapsulation was sufficient to hold the 3D mini-pillars firmly in place. For multichannel arrays, connectors (FCI/Amphenol FFC&FPC clincher connectors) were then attached by screen printing silver conductive epoxy (CircuitWorks CW2400) onto the ends of the dried MXene constructs, inserting these into the connectors, and clinching shut. The devices were baked at 70 °C for 30 min to cure the silver epoxy. Next, 1:10 PDMS was layered over the devices to form the top insulation layer, thoroughly degassed at 60 mmHg for 15 min – which forced PDMS to infiltrate into the MXene composite matrix – and subsequently cured at 70 °C for 1 h. Finally, devices were cut out with a razor blade, and peeled up from the acrylic substrate. For planar electrodes, electrode contacts were exposed by cutting circular holes though the top PDMS layer using a biopsy punch (diameters ranging from 500 μm – 3 mm) and carefully peeling up the disk of PDMS to expose the MXene composite electrode below. For 3D mini-pillar electrodes, contacts were exposed by trimming the tops of the mini-pillars with a flat razor blade, exposing the MXene-sponge composite electrode. Slight variations on this method were utilized for MXtrode arrays designed for different applications: for EMG arrays, a thin layer of silicone medical adhesive spray (Hollister Adapt 7730) was applied to the skin-facing side of the array prior to opening the electrode contacts to enhance skin adhesion; for single-channel ECG and EOG MXtrodes, EcoFlex (Smooth-on Ecoflex 00-30) in a 1:2 ratio (part A:part B) was used as the encapsulation rather than PDMS to offer enhanced skin adhesion and comfort; for the ECoG electrodes, arrays were fabricated in PDMS as described above, but were additionally coated in a 1 μm-thick layer of Parylene-C prior to opening electrode contacts to enhance the moisture barrier properties of the encapsulation.

### Imaging of MXtrode devices

Optical images of MXtrodes and their constituent parts were taken with a Keyence VHX6000 digital microscope. Scanning electron microscopy (SEM) images were captured using a Zeiss Supra 50VP scanning electron microscope with an accelerating voltage of 5 kV.

### DC conductivity of ink-infused composites

DC conductivity measurements were made on laser-cut test structures 20 cm long x 3 mm wide x 285 μm thick comprising 55% cellulose / 45% polyester blend (Texwipe TechniCloth) infused with either: (1) 30 mg/mL Ti_3_C_2_ MXene in DI, (2) 1.1% high conductivity grade PEDOT:PSS in H_2_O (Sigma Aldrich), or (3) 18 mg/mL highly concentrated single-layer graphene oxide (Graphene Supermarket) which was subsequently reduced using a vitamin-C reduction method(*98*). Measurements were taken with a handheld multimeter with flat alligator clip terminations, where the negative lead was fixed at the end of the construct and the positive lead was moved in 2 cm increments for each measurement.

### Electrochemical characterization

For cutaneous EIS measurements, the skin of the inner forearm was prepared with an alcohol swab followed by light abrasion (3M TracePrep abrasive tape) before placing 3 mm planar and 3D mini-pillar MXtrodes and measuring EIS from 1 – 10^5^ Hz with a 10 mV_pp_ driving voltage using a Gamry Reference 600 potentiostat. Reference was placed on the inner wrist and ground was placed on the elbow (Natus disposable disk electrodes). Electrochemical measurements in saline, including EIS, CV, and current pulsing, were performed for planar MXtrodes with diameters of 3 mm, 2 mm, 1 mm, and 500 μm, and Pt electrodes with a diameter of 2.3 mm (Adtec epilepsy subdural grid TG48G-SP10X-000) in 10 mM phosphate buffered saline (Quality Biological, pH 7.4) using a Gamry Reference 600 potentiostat. All electrochemical measurements were done in a 3 electrode configuration with a graphite rod counter electrode (BioRad Laboratories, Inc.) and a Ag/AgCl reference electrode (Sigma-Aldrich). EIS was measured from 1 – 10^5^ Hz with 10 mV_pp_ driving voltage. Cyclic voltammetry was performed at a sweep rate of 50 mV/s. Safe voltage limits for MXtrodes were determined by incrementally increasing the negative limit of the CV scan until water reduction was observed (beginning at −1.9 V), then the positive limit of the CV scan until a linear, resistive behavior was observed (beginning at +0.7 V) beyond which the MXtrode showed current loss with subsequent scans. CSC_c_ was determined from the CV scans by taking the time integral of the cathodal current. Current pulsing was performed using chronopotentiometry with biphasic, charge-balanced current pulses with *t_c_* = *t_a_* = 500 μs and *t_ip_* = 250 μs for currents ranging from 600 μA to 5 mA for N=3 electrodes of each size. For CIC_c_ calculations, *E_mc_* was determined as the instantaneous voltage 10 μs after the end of the cathodal current pulse. *E_mc_* values were plotted as a function of injected current amplitude, and the linear relation was determined to estimate the current limit at which the electrode would reach its cathodal limit (−1.8 V for MXtrodes, −0.6 V for Pt). Any series of measurements in which current amplitude vs. *E_mc_* was not linear with R^2^ < 0.95 were excluded. CIC_c_ was defined as: 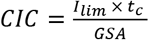, where *I_lim_* is the cathodal current limit, *t_c_* is the cathodal pulse width, and GSA is the electrode geometric surface area. CSC and CIC scaling relations were determined by fitting power functions to the data, similar to work by M. Ganji et al(*59*).

### EEG experiments

EEG experiments were conducted under a protocol approved by the Institutional Review Board of Drexel University (Protocol # 1904007140). One healthy human subject was seated in a comfortable chair with a head rest. Prior to placing electrodes, the subject’s scalp was prepared with an alcohol swab and light abrasion (3M TracePrep abrasive tape), though the presence of hair may have limited the efficacy of the skin abrasion. Recordings were made using an 8-electrode MXtrode device with dry 3 mm-diameter 3D mini-pillar electrodes and 1 standard gelled Ag/AgCl EEG cup electrode (Technomed Disposable EEG cup) placed in the center of the MXtrode array. Recordings were obtained using a NeuroNexus SmartBox amplifier system with a sampling rate of 20 kHz. Standard gelled Ag/AgCl EEG cup electrodes were used for reference (placed on left mastoid) and ground (placed on forehead center). In the first set of recordings, electrodes were positioned on the parietal region, centered over 10-20 site P1. Pre-wrap (Mueller) was used to hold the electrodes in place, and a small hole was made in this wrapping to allow application of conductive gel (SuperVisc, EASYCAP GmbH) beneath the standard EEG cup electrode. To evoke identifiable changes in the alpha band (*99*), six 2-minute-long recordings were obtained, alternating between eyes open and eyes closed states. During resting state, the subject was instructed to remain relaxed but alert. For the eyes open state, the subject was asked to fixate on a cross on a computer monitor to reduce oculomotor saccades and maintain a consistent level of external vigilance. In the second set of recordings, the hand motor region was localized using single TMS pulses to isolate a location that evoked lateral finger movements (*100*) *via* induced contractions of the first dorsal interosseus, and the electrodes were positioned centered over this location, approximately at site C3. Six 2-minute-long recordings were obtained, cycling through a resting state, imagined hand flexion, and actual hand flexion. Signals were notch filtered at 60 Hz and bandpass filtered from 0.1 – 100 Hz. Bridging analysis was performed to ensure that gel did not leak from the cup electrode to contact any recording surface on the MXtrode array.

### EMG experiments

EMG, ECG, and EOG human epidermal recordings were conducted under an experimental protocol approved by the Institutional Review Board of the University of Pennsylvania (Protocol # 831802). For all EMG experiments, skin preparation prior to placing MXtrode arrays included an alcohol swab followed by light abrasion (3M TracePrep abrasive tape), and signals were recorded at a sampling rate of 20 kHz on an Intan RHS2000 Stimulation/Recording Controller (Intan Technologies). For recordings from abductor pollicis brevis (APB), a 5×4 grid of 3 mm planar MXtrodes with center-to-center spacing of 7.5 mm horizontal, 6.5 mm vertical, was placed over the APB at the base of the thumb. Reference was placed over the bony interphalangeal thumb joint and ground was placed on the back of the hand (Natus disposable adhesive electrodes). The median nerve was stimulated at the wrist using a VikingQuest handheld bipolar stimulator (Nicolet), starting at 10 mA and gradually increasing until supramaximal activation of the APB was achieved in the form of a thumb twitch (amplitude 38.8 mA for the subject shown). For localization of the NMJ of the APB, evoked responses recorded on each electrode in the array (no signal filtering was used) were averaged across all stim trials (N=10), the peak of the average evoked response was determined, and a map of the latency of this peak from the onset of the stimulation was created. The location of the NMJ was approximated as the location with the shortest latency in the peak evoked response. For recordings of the biceps, a 10×4 grid of 3 mm planar MXtrodes with center-to-center spacing of 8.5 mm horizontal, 8.5 mm vertical, was placed over the center of the biceps muscle. Reference was placed distal to the array just above the inner elbow and ground was placed proximal to the array on the deltoid (Natus disposable adhesive electrodes). The supraclavicular nerve was stimulated using the same VikingQuest handheld bipolar stimulator (Nicolet), starting at 30 mA and gradually increasing until clear submaximal activation of biceps was observed (amplitude 49.0 mA for the subject shown). As for the APB recordings, evoked responses recorded on each electrode were averaged across all stim trials (N=11), the peak of the average evoked response was determined, and a map of the latency of this peak from the onset of the stimulation was created. The IZ was determined as the area with the shortest latency in the peak evoked response. To further validate the IZ localization in the biceps, additional EMG recordings were made with the subject performing periods of isometric voluntary contractions. Signals were subtracted in bipolar configuration down the length of the MXtrode array, and MUAPs were identified in the raw EMG signal. The IZ was determined as the region where the MUAP appears earliest and where the signal polarity inverts.

### ECG recording

For ECG recording experiments, skin preparation prior to placing electrodes included an alcohol swab followed by light abrasion (3M TracePrep abrasive tape), and signals were recorded at a 20 kHz sampling rate on an Intan RHS2000 Stimulation/Recording Controller (Intan Technologies). Recordings were made in a 3-electrode configuration with reference placed just below the subject’s right clavicle, ground placed just below the subject’s left clavicle, and the working electrode placed on the lower left ribs. For comparison, recordings were made sequentially using either all 2 cm gelled Natus electrodes (Natus disposable adhesive electrodes) or all 1.3 cm dry MXtrodes, with electrodes placed in the same locations.

### EOG recording

EOG recording experiments followed a similar protocol to ECG recording. Skin preparation prior to placing the electrodes included an alcohol swab followed by light abrasion (3M TracePrep abrasive tape), and signals were recorded at a 20 kHz sampling rate on an Intan RHS2000 Stimulation/Recording Controller (Intan Technologies). 1.3 cm-diameter dry MXtrodes were used for ground, (+) and (-) contacts. For tracking up-down eye movements, the (+) and (-) electrodes were placed below and above the right eye, respectively. The subject moved their eyes in a center-downcenter-up-center pattern for 2 minutes. For tracking leftright eye movements, the (+) and (-) electrodes were placed just lateral to the left and right eyes, respectively. The subject moved their eyes in a center-left-center-right-center pattern for 2 minutes. The ground electrode was centered on the forehead for both EOG recording paradigms.

### ECoG recording

Pigs were pair housed when possible and were always in a shared room with other pigs in a research facility certified by the Association for Assessment and Accreditation of Laboratory Animal Care International (AAALAC facility). All experiments were conducted according to the ethical guidelines set by the Institutional Animal Care and Use Committee of the University of Pennsylvania and adhered to the guidelines set forth in the NIH Public Health Service Policy on Humane Care and Use of Laboratory Animals (2015). Prior to the procedures, animals were fasted for 16 hours with water remaining ad libitum. After induction with 20 mg/kg of ketamine (Hospira, 0409–2051-05) and 0.5 mg/kg of midazolam (Hospira, 0409-2596-05), anesthesia was provided with using 2-2.5% isoflurane (Piramal, 66794-013-25) via a snout mask and glycopyrrolate was given subcutaneously to curb secretions (0.01 mg/kg; West-Ward Pharmaceutical Corp., 0143-9682-25). The animals were intubated with a size 6.0 mm endotracheal tube and anesthesia was maintained with 2–2.5% isoflurane per 2 liters O_2_. Animals were then moved to an operating room, where they were transferred onto a ventilator. The ventilator provided the same rate of isoflurane and O_2_ for anesthesia maintenance at a rate of 20-25 breaths per minute. Heart rate, respiratory rate, arterial oxygen saturation, end tidal CO_2_, blood pressure and rectal temperature were continuously monitored, while pain response to pinch was periodically assessed. All of these measures were used to maintain an adequate level of anesthesia. A forced air warming system was used to maintain normothermia. Prior to electrode insertion, the pig was placed in a stereotaxic frame described previously (*101*), and the surgical site was prepared and draped in the standard sterile fashion. After the skull was exposed, an 11 mm craniectomy was performed at the recording site, 7 mm lateral to midline and 4.5 mm posterior to bregma in order to expose the frontoparietal cortex. The dura was opened to expose the cortical surface and the MXtrode array was then placed in the subdural space for recording. Recordings were obtained using an HS-36 amplifier and collected continuously at 32 kHz using a Neuralynx Digital Lynx SX recording system. Raw data was collected and stored using Neuralynx’s Cheetah recording software.

### Cortical stimulation

An adult male Sprague Dawley rat (Crl:SD, 300 g) was used for the stimulation experiment. Anesthesia was induced with 5% isoflurane and the rat was placed in a stereotaxic frame. Depth of anesthesia was monitored by respiratory rate and pedal reflex and maintained at a surgical plane with 1.5-2.5% isoflurane. Whisker motor cortex (wM1) was exposed bilaterally with a craniotomy centered on the midline. A durotomy was performed over right wM1. A stimulation return skull screw was placed in the left frontal bone. Trains of biphasic current pulses (300 μs/phase, 3 μs interpulse interval, 100 pulses, variable pulse amplitude from 1.0 to 1.4 mA) were delivered through a single electrode in the MXtrode array using the Intan RHS System (Intan Technologies). Stimulus-evoked contralateral whisker movement was quantified with a laser micrometer (IG-028, Keyence Corp.), which had a measurement range of 28 mm, a spatial resolution of 5 μm, and a temporal resolution of 490 μs. A 360 μm diameter polyimide tube was placed over a whisker with visually apparent stimulus evoked movement. Spatial sensitivity of the micrometer was adjusted to detect only the motion of the whisker in the tube (*102*). At the conclusion of the experiment, the rat was killed with an intraperitoneal injection of sodium pentobarbital. These procedures were approved by the Institutional Animal Care and Use Committee of the University of Pennsylvania.

### Evaluation of MRI and CT compatibility

A strip of six 3 mm-diameter MXtrodes were prepared with PDMS encapsulation to match the geometry of the comparison Pt clinical ECoG electrode strips (Adtec epilepsy subdural grid TG48G-SP10X-000). Both types of electrode arrays were placed in 0.6% agarose (IBI Scientific) prepared with 10 mM PBS (Quality Biological, pH 7.4) in a 15 mm inner-diameter glass test tube, with degassing to remove air bubbles. A 9.4 T Horizontal bore MRI scanner (Bruker, Erlangen) and 35 mm diameter volume coil (m2m Imaging, USA) were used to acquire T1-weighted gradient echo MR images of cross-sections of both electrode types. The acquisition parameters for T1-W MRI were: TE/TR = 7/150 ms, FOV = 30×30 mm^2^, Matrix size = 256×256, Averages = 4, Flip angle = 30°, slice thickness = 0.7 mm. For CT imaging, a μCT50 specimen scanner (Scanco Medical, Bruttisellen, Switzerland) was used to scan the electrodes at 70 kV, 115 μA, and 10 μm isotropic resolution. 2D images in the axial plane of each electrode type were acquired for comparison.

### Magnetic susceptibility measurements

Magnetic properties of Ti_3_C_2_ were measured with Quantum Design EverCool II physical property measurement system. A free-standing film of Ti_3_C_2_ with a mass of 4.820 mg was packed in a plastic sample container. The sample was heated to 310 K and was allowed to reach thermal equilibrium for about 10 min. Magnetization was recorded with respect to the applied magnetic field up to 9 T. The measured data was subtracted from that of the plastic sample holder and normalized by sample mass.

## Acknowledgments

This work was supported by: the National Science Foundation Graduate Research Fellowship Program grant no. DGE-1845298 (N.D. and B.B.M.); the Linda Pechenik Investigator Award, Penn Health Tech Medical Device Accelerator Award, MuRata Manufacturing Co., Ltd., the Mirowski Family Foundation and Neil and Barbara Smit (F.V.); the National Institutes of Health (NIH) Office of the Director grant no. DP5-OD021352 (J.M.); the NIH training Fellowship in Neuroengineering and Medicine grant no. T32NS091006 (N.V.A.); the National Institute of Biomedical Imaging and Bioengineering (NIBIB) grant no. P41 EB015893 (R.R.); the Department of Veterans Affairs grant no. IK2 RX003118 (S.E.G.); the Department of Defense (DoD) Epilepsy Research Program (ERP) grant no. W81XWH-16-1-0675 (J.A.W.); exploration of magnetic and electrical properties of MXene was supported by U.S. Department of Energy (DOE), Office of Science, Office of Basic Energy Sciences grant no. DESC0018618 (Y.G.); Any opinions, findings, conclusions, or recommendations expressed in this material are those of the authors and do not necessarily reflect the views of the National Science Foundation. Prof. Steven J. May (Drexel University) is acknowledged for providing access to the physical property measurement system.

## Author contributions

N.D. and F.V. designed the study. N.D. and N.V.A. conceptualized the MXtrode devices. N.D. developed the fabrication process, produced all MXtrode devices, performed electrical and electrochemical characterization, captured optical microscopy images, and analyzed all of the electrophysiology data. N.D., B.E., B.B.M., conducted the EEG experiments and analyzed the data with F.V., J.M., and K.A.D. N.D., B.B.M., and G.R. performed EMG, ECG, and EOG experiments and analyzed the data with T.D. and F.V. A.G.R. performed the rat surgery and together with N.D. performed the cortical stimulation experiment. H.I.C. performed the swine surgery and together with M.S., J.A.W. and N.D. collected the ECoG data. T.M. collected SEM images and K.H. measured the magnetic properties of Ti_3_C_2_ MXene under guidance from Y.G. R.R. advised on and designed MRI experiments and P.B. collected the 9.4T MRI images. S.E.G. collected the CT images. N.D. and F.V. wrote the manuscript with contributions from all authors.

## Supplementary Materials

Fig. S1. Optical and SEM images of MXtrode composites.

Fig. S2. Scalable fabrication of MXtrode arrays.

Fig. S3. DC conductivity of ink-infused composites.

Fig. S4. Scaling of electrochemical properties for MXtrodes.

Fig. S5. EEG alpha bandpower mapping.

Fig. S6. Motor EEG recording.

Fig. S7. EMG array impedance and bipolar subtraction experiment.

Fig. S8. Electrooculography with MXtrodes.

Fig. S9. 3T MRI compatibility and magnetic susceptibility of MXene.

Table S1. Electrochemical properties of MXtrode planar electrodes of varying diameters, compared to 2.3 mm-diameter Pt electrodes and to literature values for other electrode materials.

Table S2. CSCC values for planar MXtrodes of varying diameters and the comparison clinical Pt ECoG electrode.

Movie S1. High-density dry EEG shows spatial patterns of alpha activation.

Movie S2. Cortical stimulation with MXtrodes evokes whisker movement in rat.

## References

1. J. Heikenfeld, A. Jajack, J. Rogers, P. Gutruf, L. Tian, T. Pan, R. Li, M. Khine, J. Kim, J. Wang, J. Kim, Wearable sensors: Modalities, challenges, and prospects, Lab Chip 18, 217–248 (2018).

2. J. A. Rogers, T. Someya, Y. Huang, Materials and mechanics for stretchable electronics, Science (80-.). 327, 1603–1607 (2010).

3. Y. Liu, M. Pharr, G. A. Salvatore, Lab-on-Skin: A Review of Flexible and Stretchable Electronics for Wearable Health Monitoring, ACS Nano 11, 9614–9635 (2017).

4. F. Vitale, B. Litt, Bioelectronics: the promise of leveraging the body’s circuitry to treat disease, Bioelectron. Med. 1, 3–7 (2018).

5. J. J. S. Norton, D. S. Lee, J. W. Lee, W. Lee, O. Kwon, P. Won, S. Y. Jung, H. Cheng, J. W. Jeong, A. Akce, S. Umunna, I. Na, Y. H. Kwon, X. Q. Wang, Z. J. Liu, U. Paik, Y. Huang, T. Bretl, W. H. Yeo, J. A. Rogers, Z. Bao, Soft, curved electrode systems capable of integration on the auricle as a persistent brain-computer interface, Proc. Natl. Acad. Sci. U. S. A. 112, 3920–3925 (2015).

6. J. W. Jeong, W. H. Yeo, A. Akhtar, J. J. S. Norton, Y. J. Kwack, S. Li, S. Y. Jung, Y. Su, W. Lee, J. Xia, H. Cheng, Y. Huang, W. S. Choi, T. Bretl, J. A. Rogers, Materials and optimized designs for human-machine interfaces via epidermal electronics, Adv. Mater. 25, 6839–6846 (2013).

7. B. Xu, A. Akhtar, Y. Liu, H. Chen, W. H. Yeo, S. Park, B. Boyce, H. Kim, J. Yu, H. Y. Lai, S. Jung, Y. Zhou, J. Kim, S. Cho, Y. Huang, T. Bretl, J. A. Rogers, An Epidermal Stimulation and Sensing Platform for Sensorimotor Prosthetic Control, Management of Lower Back Exertion, and Electrical Muscle Activation, Adv. Mater. 28, 4462–4471 (2016).

8. D. J. McFarland, J. R. Wolpaw, EEG-based brain–computer interfaces, Curr. Opin. Biomed. Eng. 4, 194–200 (2017).

9. S. Lee, W. Y. X. Peh, J. Wang, F. Yang, J. S. Ho, N. V. Thakor, S. C. Yen, C. Lee, Toward Bioelectronic Medicine—Neuromodulation of Small Peripheral Nerves Using Flexible Neural Clip, Adv. Sci. 4, 1700149 (2017).

10. M. R. DeLong, T. Wichmann, Basal ganglia circuits as targets for neuromodulation in Parkinson disease JAMA Neurol. 72, 1354–1360 (2015).

11. H. Irisawa, H. F. Brown, W. Giles, Cardiac pacemaking in the sinoatrial nodePhysiol. Rev. 73, 197–227 (1993).

12. G. T. Hwang, H. Park, J. H. Lee, S. Oh, K. Il Park, M. Byun, H. Park, G. Ahn, C. K. Jeong, K. No, H. Kwon, S. G. Lee, B. Joung, K. J. Lee, Self-powered cardiac pacemaker enabled by flexible single crystalline PMN-PT piezoelectric energy harvester, Adv. Mater. 26, 4880–4887 (2014).

13. L. Ulloa, S. Quiroz-Gonzalez, R. Torres-Rosas, Nerve Stimulation: Immunomodulation and Control of InflammationTrends Mol. Med. 23, 1103–1120 (2017).

14. J. Cheng, H. Shen, R. Chowdhury, T. Abdi, F. Selaru, J. D. Z. Chen, Potential of Electrical Neuromodulation for Inflammatory Bowel Disease, Inflamm. Bowel Dis. 26, 1119–1130 (2020).

15. M. H. Granat, A. C. B. Ferguson, B. J. Andrews, M. Delargy, The role of functional electrical stimulation in the rehabilitation of patients with incomplete spinal cord injury observed benefits during gait studies, Paraplegia 31, 207–215 (1993).

16. P. H. Peckham, J. S. Knutson, Functional Electrical Stimulation for Neuromuscular Applications, Annu. Rev. Biomed. Eng. 7, 327–360 (2005).

17. C. Ethier, E. R. Oby, M. J. Bauman, L. E. Miller, Restoration of grasp following paralysis through brain-controlled stimulation of muscles, Nature 485, 368–371 (2012).

18. R. A. Nawrocki, H. Jin, S. Lee, T. Yokota, M. Sekino, T. Someya, Self-Adhesive and Ultra-Conformable, Sub-300 nm Dry Thin-Film Electrodes for Surface Monitoring of Biopotentials, Adv. Funct. Mater. 28, 1803279 (2018).

19. L. Tian, B. Zimmerman, A. Akhtar, K. J. Yu, M. Moore, J. Wu, R. J. Larsen, J. W. Lee, J. Li, Y. Liu, B. Metzger, S. Qu, X. Guo, K. E. Mathewson, J. A. Fan, J. Cornman, M. Fatina, Z. Xie, Y. Ma, J. Zhang, Y. Zhang, F. Dolcos, M. Fabiani, G. Gratton, T. Bretl, L. J. Hargrove, P. V. Braun, Y. Huang, J. A. Rogers, Large-area MRI-compatible epidermal electronic interfaces for prosthetic control and cognitive monitoring, Nat. Biomed. Eng. 3, 194–205 (2019).

20. D. H. Kim, N. Lu, R. Ma, Y. S. Kim, R. H. Kim, S. Wang, J. Wu, S. M. Won, H. Tao, A. Islam, K. J. Yu, T. Il Kim, R. Chowdhury, M. Ying, L. Xu, M. Li, H. J. Chung, H. Keum, M. McCormick, P. Liu, Y. W. Zhang, F. G. Omenetto, Y. Huang, T. Coleman, J. A. Rogers, Epidermal electronics, Science (80-.). 333, 838–843 (2011).

21. J. Viventi, D.-H. Kim, L. Vigeland, E. S. Frechette, J. a Blanco, Y.-S. Kim, A. E. Avrin, V. R. Tiruvadi, S.-W. Hwang, A. C. Vanleer, D. F. Wulsin, K. Davis, C. E. Gelber, L. Palmer, J. Van der Spiegel, J. Wu, J. Xiao, Y. Huang, D. Contreras, J. A. Rogers, B. Litt, Flexible, foldable, actively multiplexed, high-density electrode array for mapping brain activity in vivo, Nat. Neurosci. 14, 1599–605 (2011).

22. D. Khodagholy, T. Doublet, M. Gurfinkel, P. Quilichini, E. Ismailova, P. Leleux, T. Herve, S. Sanaur, C. Bernard, G. F. Malliaras, Highly conformable conducting polymer electrodes for in vivo recordings, Adv. Mater. 23, H268–H272 (2011).

23. J. Li, P. Wang, H. J. Huang, Dry Epidermal Electrodes Can Provide Long-Term High Fidelity Electromyography for Limited Dynamic Lower Limb Movements, Sensors (Basel). 20, 4848 (2020).

24. A. Searle, L. Kirkup, A direct comparison of wet, dry and insulating bioelectric recording electrodes, Physiol. Meas. 21, 271–283 (2000).

25. G. C. Jensen, C. E. Krause, G. A. Sotzing, J. F. Rusling, Inkjet-printed gold nanoparticle electrochemical arrays on plastic. Application to immunodetection of a cancer biomarker protein, Phys. Chem. Chem. Phys. 13, 4888–4894 (2011).

26. G. Mattana, A. Loi, M. Woytasik, M. Barbaro, V. Noël, B. Piro, Inkjet-Printing: A New Fabrication Technology for Organic TransistorsAdv. Mater. Technol. 2, 1700063 (2017).

27. N. Lewinski, V. Colvin, R. Drezek, Cytotoxicity of nanopartides Small 4, 26–49 (2008).

28. M. Gao, L. Li, Y. Song, Inkjet printing wearable electronic devices J. Mater. Chem. C 5, 2971–2993 (2017).

29. U. Kraft, F. Molina-Lopez, D. Son, Z. Bao, B. Murmann, Ink Development and Printing of Conducting Polymers for Intrinsically Stretchable Interconnects and Circuits, Adv. Electron. Mater. 6, 1900681 (2020).

30. S. Park, J. An, J. R. Potts, A. Velamakanni, S. Murali, R. S. Ruoff, Hydrazine-reduction of graphite-and graphene oxide, Carbon N. Y. 49, 3019–3023 (2011).

31. C. Mattevi, G. Eda, S. Agnoli, S. Miller, K. A. Mkhoyan, O. Celik, D. Mastrogiovanni, G. Granozzi, E. Carfunkel, M. Chhowalla, Evolution of electrical, chemical, and structural properties of transparent and conducting chemically derived graphene thin films, Adv. Funct. Mater. 19, 2577–2583 (2009).

32. K. Hantanasirisakul, M. Q. Zhao, P. Urbankowski, J. Halim, B. Anasori, S. Kota, C. E. Ren, M. W. Barsoum, Y. Gogotsi, Fabrication of Ti_3_C_2_ MXene Transparent Thin Films with Tunable Optoelectronic Properties, Adv. Electron. Mater. 2, 1600050 (2016).

33. M. Mariano, O. Mashtalir, F. Q. Antonio, W. H. Ryu, B. Deng, F. Xia, Y. Gogotsi, A. D. Taylor, Solution-processed titanium carbide MXene films examined as highly transparent conductors, Nanoscale 8, 16371–16378 (2016).

34. S. J. Kim, J. Choi, K. Maleski, K. Hantanasirisakul, H. T. Jung, Y. Gogotsi, C. W. Ahn, Interfacial assembly of ultrathin, functional MXene films, ACS Appl. Mater. Interfaces 11, 32320–32327 (2019).

35. E. Quain, T. S. Mathis, N. Kurra, K. Maleski, K. L. Van Aken, M. Alhabeb, H. N. Alshareef, Y. Gogotsi, Direct Writing of Additive-Free MXene-in-Water Ink for Electronics and Energy Storage, Adv. Mater. Technol. 4, 1800256 (2019).

36. C. Zhang, L. McKeon, M. P. Kremer, S.-H. Park, O. Ronan, A. Seral-Ascaso, S. Barwich, C. Ó. Coileáin, N. McEvoy, H. C. Nerl, B. Anasori, J. N. Coleman, Y. Gogotsi, V. Nicolosi, Additive-free MXene inks and direct printing of micro-supercapacitors, Nat. Commun. 10, 1795 (2019).

37. C. Dai, H. Lin, G. Xu, Z. Liu, R. Wu, Y. Chen, Biocompatible 2D Titanium Carbide (MXenes) Composite Nanosheets for pH-Responsive MRI-Guided Tumor Hyperthermia, Chem. Mater. 29, 8637–8652 (2017).

38. F. Vitale, N. Driscoll, B. Murphy, in 2D Metal Carbides and Nitrides (MXenes): Structure, Properties and Applications, B. Anasori, Y. Gogotsi, Eds. (Springer International Publishing, Cham, Cham, 2019), pp. 503–524.

39. X. Han, J. Huang, H. Lin, Z. Wang, P. Li, Y. Chen, 2D Ultrathin MXene-Based Drug-Delivery Nanoplatform for Synergistic Photothermal Ablation and Chemotherapy of Cancer, Adv. Healthc. Mater. 7, 1701394 (2018).

40. F. Meng, M. Seredych, C. Chen, V. Gura, S. Mikhalovsky, S. Sandeman, G. Ingavle, T. Ozulumba, L. Miao, B. Anasori, Y. Gogotsi, MXene Sorbents for Removal of Urea from Dialysate: A Step toward the Wearable Artificial Kidney, ACS Nano 12, 10518–10528 (2018).

41. L. Li, X. Fu, S. Chen, S. Uzun, A. S. Levitt, C. E. Shuck, W. Han, Y. Gogotsi, Hydrophobic and Stable MXene-Polymer Pressure Sensors for Wearable Electronics, Cite This ACS Appl. Mater. Interfaces 12, 15362–15369 (2020).

42. S. Seyedin, S. Uzun, A. Levitt, B. Anasori, G. Dion, Y. Gogotsi, J. M. Razal, MXene Composite and Coaxial Fibers with High Stretchability and Conductivity for Wearable Strain Sensing Textiles, Adv. Funct. Mater. 30, 1910504 (2020).

43. N. Driscoll, A. G. Richardson, K. Maleski, B. Anasori, O. Adewole, P. Lelyukh, L. Escobedo, D. K. Cullen, T. H. Lucas, Y. Gogotsi, F. Vitale, Two-Dimensional Ti_3_C_2_MXene for High-Resolution Neural Interfaces, ACS Nano 12, 10419–10429 (2018).

44. N. Driscoll, K. Maleski, A. G. Richardson, B. Murphy, B. Anasori, T. H. Lucas, Y. Gogotsi, F. Vitale, Fabrication of Ti_3_C_2_ MXene Microelectrode Arrays for In Vivo Neural Recording, J. Vis. Exp. 15, 1–9 (2020).

45. B. B. Murphy, P. J. Mulcahey, N. Driscoll, A. G. Richardson, G. T. Robbins, N. V Apollo, K. Maleski, T. H. Lucas, Y. Gogotsi, T. Dillingham, F. Vitale, A Gel-Free Ti_3_C_2_T_x_-Based Electrode Array for High-Density, High-Resolution Surface Electromyography, Adv. Mater. Technol. 5, 1–10 (2020).

46. J. Zhang, N. Kong, S. Uzun, A. Levitt, S. Seyedin, P. A. Lynch, S. Qin, M. Han, W. Yang, J. Liu, X. Wang, Y. Gogotsi, J. M. Razal, Scalable Manufacturing of Free-Standing, Strong Ti_3_C_2_T_x_ MXene Films with Outstanding Conductivity, Adv. Mater. 32(2020), doi: 10.1002/adma.202001093.

47. T. Mathis, K. Maleski, A. Goad, A. Sarycheva, M. Anayee, A. C. Foucher, E. Stach, Y. Gogotsi, C. Mathis, T. S. Mathis, K. Maleski, A. Goad, A. Sarycheva, M. Anayee, C. Alexandre, Modified MAX Phase Synthesis for Environmentally Stable and Highly Conductive Ti_3_C_2_MXene Modified MAX Phase Synthesis for Environmentally Stable and Highly Conductive Ti_3_C_2_MXene, ChemRxiv (2020), doi: 10.26434/chemrxiv._12805280.v1.

48. C. E. Shuck, A. Sarycheva, M. Anayee, A. Levitt, Y. Zhu, S. Uzun, V. Balitskiy, V. Zahorodna, O. Gogotsi, Y. Gogotsi, Scalable Synthesis of Ti_3_C_2_T_x_ MXene, Adv. Eng. Mater. 22, 1901241 (2020).

49. M. Alhabeb, K. Maleski, B. Anasori, P. Lelyukh, L. Clark, S. Sin, Y. Gogotsi, Guidelines for Synthesis and Processing of Two-Dimensional Titanium Carbide (Ti_3_C_2_T_x_ MXene), Chem. Mater. 29, 7633–7644 (2017).

50. S. Uzun, S. Seyedin, A. L. Stoltzfus, A. S. Levitt, M. Alhabeb, M. Anayee, C. J. Strobel, J. M. Razal, G. Dion, Y. Gogotsi, Knittable and Washable Multifunctional MXene-Coated Cellulose Yarns, Adv. Funct. Mater. 29, 1905015 (2019).

51. W. Franks, I. Schenker, P. Schmutz, A. Hierlemann, Impedance characterization and modeling of electrodes for biomedical applications, IEEE Trans. Biomed. Eng. 52, 1295–1302 (2005).

52. S. F. Cogan, P. R. Troyk, J. Ehrlich, T. D. Plante, D. E. Detlefsen, Potential-biased, asymmetric waveforms for charge-injection with activated iridium oxide (AIROF) neural stimulation electrodes, IEEE Trans. Biomed. Eng. 53, 327–332 (2006).

53. W. M. Grill, J. T. Mortimer, Stimulus Waveforms for Selective Neural Stimulation, IEEE Eng. Med. Biol. Mag. 14, 375–385 (1995).

54. S. F. Cogan, Neural stimulation and recording electrodes., Annu. Rev. Biomed. Eng. 10, 275–309 (2008).

55. L. Lorencova, T. Bertok, J. Filip, M. Jerigova, D. Velic, P. Kasak, K. A. Mahmoud, J. Tkac, Highly stable Ti_3_C_2_T_x_ MXene)/Pt nanoparticles-modified glassy carbon electrode for H2O2 and small molecules sensing applications, Sensors Actuators, B Chem. 263, 360–368 (2018).

56. Z. Ling, C. E. Ren, M. Q. Zhao, J. Yang, J. M. Giammarco, J. Qiu, M. W. Barsoum, Y. Gogotsi, Flexible and conductive MXene films and nanocomposites with high capacitance, Proc. Natl. Acad. Sci. U. S. A. 111, 16676–16681 (2014).

57. C. J. Zhang, B. Anasori, A. Seral-Ascaso, S.-H. Park, N. McEvoy, A. Shmeliov, G. S. Duesberg, J. N. Coleman, Y. Gogotsi, V. Nicolosi, Transparent, Flexible, and Conductive 2D Titanium Carbide (MXene) Films with High Volumetric Capacitance, Adv. Mater. 29, 1702678 (2017).

58. B. Anasori, M. R. Lukatskaya, Y. Gogotsi, 2D metal carbides and nitrides (MXenes) for energy storage, Nat. Rev. Mater. 2, 16098 (2017).

59. M. Ganji, A. Tanaka, V. Gilja, E. Halgren, S. A. Dayeh, Scaling Effects on the Electrochemical Stimulation Performance of Au, Pt, and PEDOT:PSS Electrocorticography Arrays, Adv. Funct. Mater. 27, 1703019 (2017).

60. D. J. Hewson, Y. Langeron, J. Duche, Evolution in impedance at the electrode-skin interface of two types of surface EMG electrodes during long-term recordings, J. Electromyogr. Kinesiol. 13, 273–279 (2003).

61. G. Li, S. Wang, Y. Y. Duan, Towards gel-free electrodes: A systematic study of electrode-skin impedance, Sensors Actuators, B Chem. 241, 1244–1255 (2017).

62. S. M. Lee, H. J. Byeon, J. H. Lee, D. H. Baek, K. H. Lee, J. S. Hong, S. H. Lee, Self-adhesive epidermal carbon nanotube electronics for tether-free long-term continuous recording of biosignals, Sci. Rep. 4, 6074 (2014).

63. Y. H. Chen, M. Op de Beeck, L. Vanderheyden, E. Carrette, V. Mihajlović, K. Vanstreels, B. Grundlehner, S. Gadeyne, P. Boon, C. van Hoof, Soft, comfortable polymer dry electrodes for high quality ECG and EEG recording, Sensors 14, 23758–23780 (2014).

64. T. C. Ferree, P. Luu, G. S. Russell, D. M. Tucker, Scalp electrode impedance, infection risk, and EEG data quality, Clin. Neurophysiol. 112, 536–544 (2001).

65. E. S. Kappenman, S. J. Luck, The effects of electrode impedance on data quality and statistical significance in ERP recordings, Psychophysiology 47, 888–904 (2010).

66. W. Klimesch, EEG alpha and theta oscillations reflect cognitive and memory performance: A review and analysisBrain Res. Rev. 29, 169–195 (1999).

67. S. W. Hughes, V. Crunelli, Thalamic mechanisms of EEG alpha rhythms and their pathological implications, Neuroscientist 11, 357–372 (2005).

68. Y. Li, J. Long, T. Yu, Z. Yu, C. Wang, H. Zhang, C. Guan, An EEG-based BCI system for 2-D cursor control by combining Mu/Beta rhythm and P300 potential, IEEE Trans. Biomed. Eng. 57, 2495–2505 (2010).

69. H.-J. Hwang, K. Kwon, C.-H. Im, Neurofeedback-based motor imagery training for brain-computer interface (BCI), J. Neurosci. Methods 179, 150–156 (2009).

70. G. Drost, D. F. Stegeman, B. G. M. van Engelen, M. J. Zwarts, Clinical applications of high-density surface EMG: A systematic review. J. Electromyogr. Kinesiol. 16, 586–602 (2006).

71. E. Martinez-Valdes, C. M. Laine, D. Falla, F. Mayer, D. Farina, High-density surface electromyography provides reliable estimates of motor unit behavior, Clin. Neurophysiol. 127, 2534–2541 (2016).

72. E. Chang, N. Ghosh, D. Yanni, S. Lee, D. Alexandru, T. Mozaffar, A Review of Spasticity Treatments: Pharmacological and Interventional Approaches, Crit. Rev. Phys. Rehabil. Med. 25, 11–22 (2013).

73. A. P. Yelnik, O. Simon, B. Parratte, J. M. Gracies, How to clinically assess and treat muscle overactivity in spastic paresis. J. Rehabil. Med. 42, 801–807 (2010).

74. S. Wynter, L. Dissabandara, A comprehensive review of motor innervation of the hand: variations and clinical significance, Surg. Radiol. Anat. 40, 259–269 (2018).

75. T. Masuda, T. Sadoyama, Distribution of innervation zones in the human biceps brachii, J. Electromyogr. Kinesiol. 1, 107–115 (1991).

76. T. L. Morris, J. C. Miller, Electrooculographic and performance indices of fatigue during simulated flight, Biol. Psychol. 42, 343–360 (1996).

77. A. B. Usakli, S. Gurkan, F. Aloise, G. Vecchiato, F. Babiloni, On the use of electrooculogram for efficient human computer interfaces, Comput. Intell. Neurosci. 2010, 135629 (2010).

78. J. I. Berman, M. S. Berger, P. Mukherjee, R. G. Henry, Diffusion-tensor imaging-guided tracking of fibers of the pyramidal tract combined with intraoperative cortical stimulation mapping in patients with gliomas, J. Neurosurg. 101, 66–72 (2004).

79. M. J. Morrell, Responsive cortical stimulation for the treatment of medically intractable partial epilepsy, Neurology 77, 1295–1304 (2011).

80. B. Lee, D. Kramer, M. A. Salas, S. Kellis, D. Brown, T. Dobreva, C. Klaes, C. Heck, C. Liu, R. A. Andersen, Engineering artificial somatosensation through cortical stimulation in humans, Front. Syst. Neurosci. 12, 24 (2018).

81. J. F. Schenck, The role of magnetic susceptibility in magnetic resonance imaging: MRI magnetic compatibility of the first and second kinds, Med. Phys. 23, 815–850 (1996).

82. J. H. Duyn, The future of ultra-high field MRI and fMRI for study of the human brainNeuroimage 62, 1241–1248 (2012).

83. K. A. Davis, R. P. R. Nanga, S. Das, S. H. Chen, P. N. Hadar, J. R. Pollard, T. H. Lucas, R. T. Shinohara, B. Litt, H. Hariharan, M. A. Elliott, J. A. Detre, R. Reddy, Glutamate imaging (GluCEST) lateralizes epileptic foci in nonlesional temporal lobe epilepsy, Sci. Transl. Med. 7, 309ra161 (2015).

84. H. Wei, O. T. Bruns, M. G. Kaul, E. C. Hansen, M. Barch, A. Wisniowska, O. Chen, Y. Chen, N. Li, S. Okada, J. M. Cordero, M. Heine, C. T. Farrar, D. M. Montana, G. Adam, H. Ittrich, A. Jasanoff, P. Nielsen, M. G. Bawendi, Exceedingly small iron oxide nanoparticles as positive MRI contrast agents, Proc. Natl. Acad. Sci. U. S. A. 114, 2325–2330 (2017).

85. W. Neumann, T. P. Pusch, M. Siegfarth, L. R. Schad, J. L. Stallkamp, CT and MRI compatibility of flexible 3D-printed materials for soft actuators and robots used in image-guided interventions, Med. Phys. 46, 5488–5498 (2019).

86. Z. Fan, Y. Wang, Z. Xie, D. Wang, Y. Yuan, H. Kang, B. Su, Z. Cheng, Y. Liu, Modified MXene/Holey Graphene Films for Advanced Supercapacitor Electrodes with Superior Energy Storage, Adv. Sci. 5, 1800750 (2018).

87. H. Daley, K. Englehart, L. Hargrove, U. Kuruganti, High density electromyography data of normally limbed and transradial amputee subjects for multifunction prosthetic control, J. Electromyogr. Kinesiol. 22, 478–484 (2012).

88. D. Farina, T. Lorrain, F. Negro, N. Jiang, High-density EMG E-Textile systems for the control of active prostheses, 32nd Annu. Int. Conf. IEEE Eng. Med. Biol., 3591–3593 (2010).

89. M. Murphy, M. A. Bruno, B. A. Riedner, P. Boveroux, Q. Noirhomme, E. C. Landsness, J. F. Brichant, C. Phillips, M. Massimini, S. Laureys, G. Tononi, M. Boly, Propofol anesthesia and sleep: A high-density EEG study, Sleep 34, 283–291 (2011).

90. M. Massimini, G. Tononi, R. Huber, Slow waves, synaptic plasticity and information processing: insights from transcranial magnetic stimulation and high-density EEG experiments, Eur. J. Neurosci. 29, 1761–1770 (2009).

91. C. Guger, H. Ramoser, G. Pfurtscheller, Real-time EEG analysis with subject-specific spatial patterns for a brain-computer interface (BCI), IEEE Trans. Rehabil. Eng. 8, 447–456 (2000).

92. B. Blankertz, K. R. Müller, G. Curio, T. M. Vaughan, G. Schalk, J. R. Wolpaw, A. Schlögl, C. Neuper, G. Pfurtscheller, T. Hinterberger, M. Schröder, N. Birbaumer, The BCI competition 2003: Progress and perspectives in detection and discrimination of EEG single trialsIEEE Trans. Biomed. Eng. 51, 1044–1051 (2004).

93. M. Jochumsen, I. K. Niazi, K. Dremstrup, E. N. Kamavuako, Detecting and classifying three different hand movement types through electroencephalography recordings for neurorehabilitation, Med. Biol. Eng. Comput. 54, 1491–1501 (2016).

94. M. Van Vliet, A. Robben, N. Chumerin, N. V. Manyakov, A. Combaz, M. M. Van Hulle, in 2012 ISSNIP Biosignals and Biorobotics Conference: Biosignals and Robotics for Better and Safer Living (BRC), (2012), pp. 1–6.

95. K. Cai, M. Haris, A. Singh, F. Kogan, J. H. Greenberg, G. Hariharan, J. A. Detre, R. Reddy, Magnetic resonance imaging of glutamate, Nat. Med. 18, 302–306 (2012).

96. F. Shahzad, A. Iqbal, S. A. Zaidi, S. W. Hwang, C. M. Koo, Nafion-stabilized two-dimensional transition metal carbide (Ti_3_C_2_T_x_ MXene) as a high-performance electrochemical sensor for neurotransmitter, J. Ind. Eng. Chem. 79, 338–344 (2019).

97. X. Zhao, A. Vashisth, E. Prehn, W. Sun, S. A. Shah, T. Habib, Y. Chen, Z. Tan, J. L. Lutkenhaus, M. Radovic, M. J. Green, Antioxidants Unlock Shelf-Stable Ti_3_C_2_T_x_ (MXene) Nanosheet Dispersions, Matter 1, 513–526 (2019).

98. J. Zhang, H. Yang, G. Shen, P. Cheng, J. Zhang, S. Guo, Reduction of graphene oxide via l-ascorbic acid, Chem. Commun. 46, 1112–1114 (2010).

99. R. J. Barry, A. R. Clarke, S. J. Johnstone, C. A. Magee, J. A. Rushby, EEG differences between eyes-closed and eyes-open resting conditions, Clin. Neurophysiol. 118, 2765–2773 (2007).

100. Z. D. Jonker, R. van der Vliet, C. M. Hauwert, C. Gaiser, J. H. M. Tulen, J. N. van der Geest, O. Donchin, G. M. Ribbers, M. A. Frens, R. W. Selles, TMS motor mapping: Comparing the absolute reliability of digital reconstruction methods to the golden standard, Brain Stimul. 12, 309–313 (2019).

101. A. V. Ulyanova, P. F. Koch, C. Cottone, M. R. Grovola, C. D. Adam, K. D. Browne, M. T. Weber, R. J. Russo, K. G. Gagnon, D. H. Smith, H. Isaac Chen, V. E. Johnson, D. Kacy Cullen, J. A. Wolf, Electrophysiological signature reveals laminar structure of the porcine hippocampus, eNeuro 5, e0102–18.2018 (2018).

102. M. A. Attiah, J. de Vries, A. G. Richardson, T. H. Lucas, A Rodent Model of Dynamic Facial Reanimation Using Functional Electrical Stimulation, Front. Neurosci. 11, 193 (2017).

103. M. Ganji, A. T. Elthakeb, A. Tanaka, V. Gilja, E. Halgren, S. A. Dayeh, Scaling Effects on the Electrochemical Performance of poly(3,4-ethylenedioxythiophene (PEDOT), Au, and Pt for Electrocorticography Recording, Adv. Funct. Mater. 27, 1703018 (2017).

104. R. A. Green, H. Toor, C. Dodds, N. H. Lovell, Variation in performance of platinum electrodes with size and surface roughness, Sensors Mater. 24, 165–180 (2012).

105. X. Luo, C. L. Weaver, D. D. Zhou, R. Greenberg, X. T. Cui, Highly stable carbon nanotube doped poly(3,4-ethylenedioxythiophene) for chronic neural stimulation, Biomaterials 32, 5551–5557 (2011).

106. Y. Lu, H. Lyu, A. G. Richardson, T. H. Lucas, D. Kuzum, Flexible Neural Electrode Array Based-on Porous Graphene for Cortical Microstimulation and Sensing, Sci. Rep. 6, 33526 (2016).

107. F. Vitale, S. R. Summerson, B. Aazhang, C. Kemere, M. Pasquali, Neural Stimulation and Recording with Bidirectional, Soft Carbon Nanotube Fiber Microelectrodes, ACS Nano 9, 4465–4474 (2015).

108. J. D. Weiland, D. J. Anderson, M. S. Humayun, In vitro electrical properties for iridium oxide versus titanium nitride stimulating electrodes, IEEE Trans. Biomed. Eng. 49, 1574–1579 (2002).

109. C. Jiang, L. Li, H. Hao, Carbon nanotube yarns for deep brain stimulation electrode, IEEE Trans. Neural Syst. Rehabil. Eng. 19, 612–616 (2011).

