## Supplementary Information for "MXtrodes: MXene-infused bioelectronic interfaces for multiscale electrophysiology and stimulation"

**Supplementary Materials**

Fig. S1. Optical and SEM images of MXtrode composites.

Fig. S2. Scalable fabrication of MXtrode arrays.

Fig. S3. DC conductivity of ink-infused composites.

Fig. S4. Scaling of electrochemical properties for MXtrodes.

Fig. S5. EEG alpha bandpower mapping.

Fig. S6. Motor EEG recording.

Fig. S7. EMG array impedance and bipolar subtraction experiment.

Fig. S8. Electrooculography with MXtrodes.

Fig. S9. 3T MRI compatibility and magnetic susceptibility of MXene.

Table S1. Electrochemical properties of MXtrode planar electrodes of varying diameters, compared to 2.3 mm-diameter Pt electrodes and to literature values for other electrode materials.

Table S2. CSCC values for planar MXtrodes of varying diameters and the comparison clinical Pt ECoG electrode.

Movie S1. High-density dry EEG shows spatial patterns of alpha activation.

Movie S2. Cortical stimulation with MXtrodes evokes whisker movement in rat.

[1]Department of Bioengineering, University of Pennsylvania, Philadelphia, PA, USA, [2]Center for Neuroengineering and Therapeutics, University of Pennsylvania, Philadelphia, PA, USA, [3]Center for Neurotrauma, Neurodegeneration, and Restoration, Corporal Michael J. Crescenz Veterans Affairs Medical Center, Philadelphia, PA, USA, [4]Department of Psychology, Drexel University, Philadelphia, PA, USA, [5]Department of Neurosurgery, University of Pennsylvania, Philadelphia, PA, USA, [6]Department of Physical Medicine and Rehabilitation, University of Pennsylvania, PA, USA, [7]Department of Materials Science and Engineering, Drexel University, Philadelphia, PA, USA, [8]A.J. Drexel Nanomaterials Institute, Drexel University, Philadelphia, PA, USA, [9]Department of Radiology, Center for Magnetic Resonance and Optical Imaging, University of Pennsylvania, Philadelphia, PA, USA, [10]Diagnostic Imaging, St Jude Children's Research Hospital, Memphis, TN, USA, [11]Translational Musculoskeletal Research Center, Corporal Michael J. Crescenz VA Medical Center, Philadelphia, PA, USA, [12]McKay Orthopaedic Research Laboratory, Department of Orthopaedic Surgery, University of Pennsylvania, Philadelphia, PA, USA, [13]Department of Neurology, University of Pennsylvania, Philadelphia, PA, USA, [14]Department of Neurology, Drexel University, Philadelphia, PA, USA

2020

### Supplementary Figures

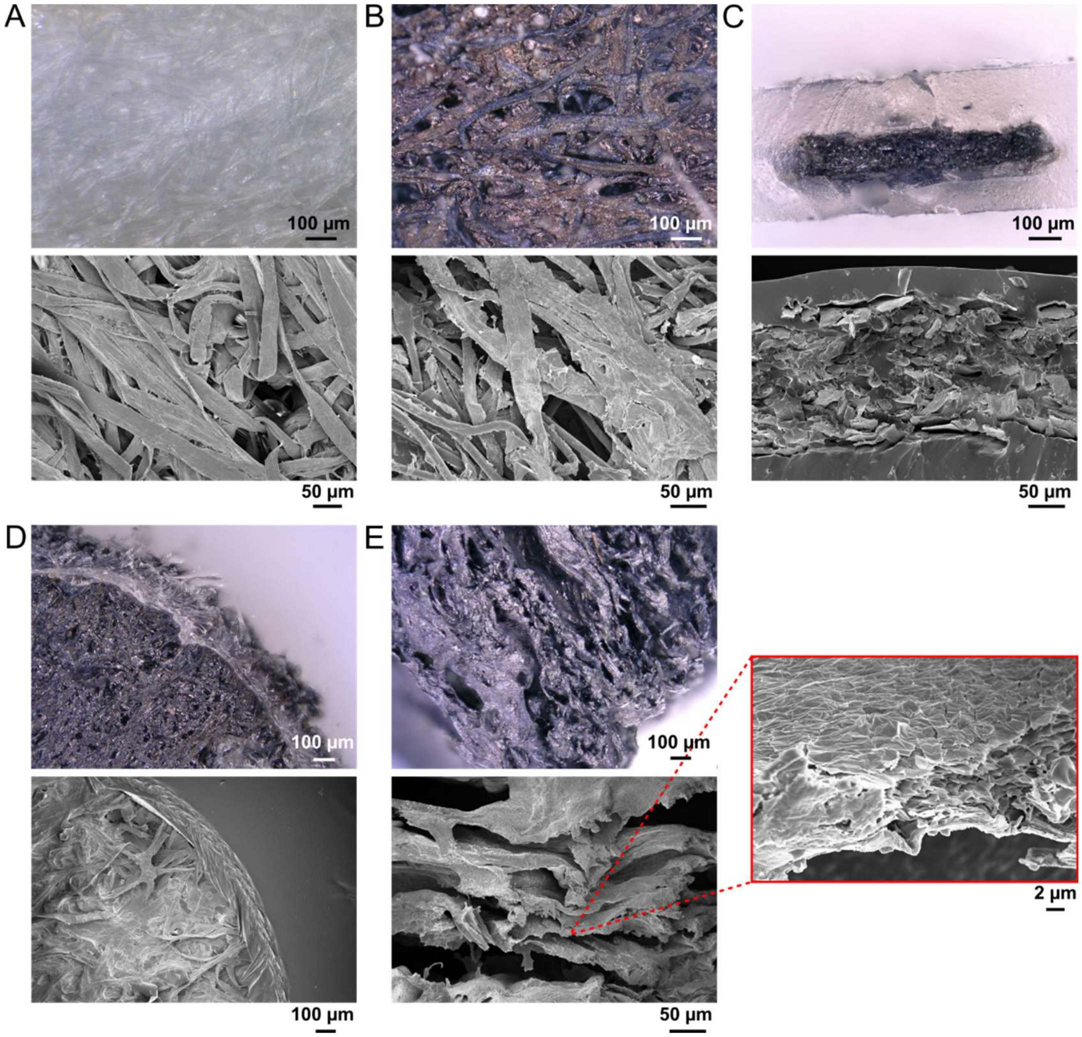


**Fig. S1. Optical and SEM images of MXtrode composites**. Optical microscopy images (top panel) and corresponding SEM images (bottom panel) for: (**A**) pristine cellulose/polyester blend substrate, (**B**) the same substrate after infusing with MXene ink, (**C**) cross-section of MXene composite trace embedded in PDMS, (**D**) edge of planar electrode contact, (**E**) side of MXene-infused cellulose foam in 3D mini-pillar MXtrode.

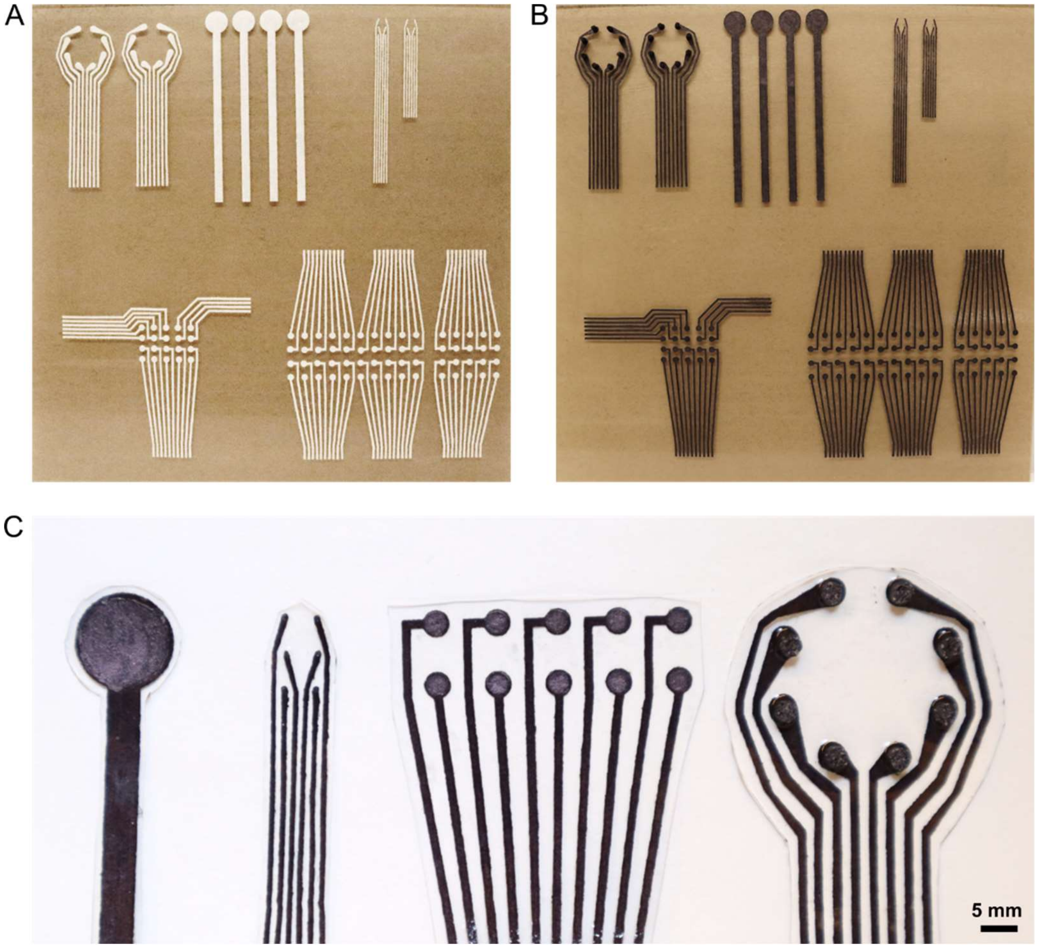


**Fig. S2. Scalable fabrication of MXtrode arrays**. (**A**) Photo of laser-patterned array substrates for various device and array geometries. (**B**) The same batch of devices shown in (A) after infiltrating with MXene ink. In the top left are shown the EEG ring MXtrode arrays after addition of the 3D mini-pillars. (**C**) Photographs of completed devices (from left to right) designed for ECG, ECoG, EMG, and EEG sensing.

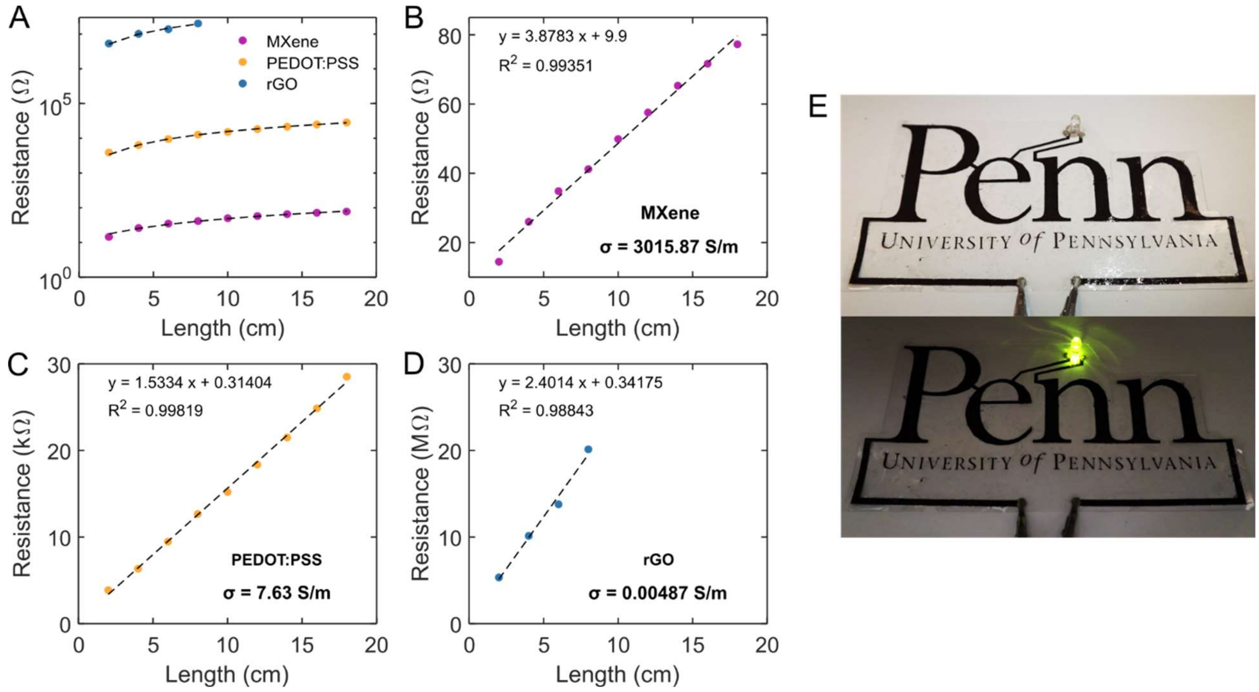


**Fig. S3. DC conductivity of ink-infused composites.** (**A**) Plot of length vs. DC resistance for composites made using MXene, PEDOT:PSS, and rGO as the conductive ink. Test structures were 20 cm x 3 mm x 285 μm (L x W x H) strips. (**B-D**) Individual plots of DC resistance *vs.* length for (**B**) MXene, (**C**) PEDOT:PSS, and (**D**) rGO, with linear fitting curves shown as dashed lines. The linear relation of resistance *vs.* length, along with the cross-sectional area of the test structure, is used to compute the bulk conductivity, **σ**, of the composites. DC resistance of the rGO composite could only be measured out to 8 cm due to high resistance. (**E**) Demonstration of exceptional conductivity of the MXene composite here used as conductive trace to power an LED.

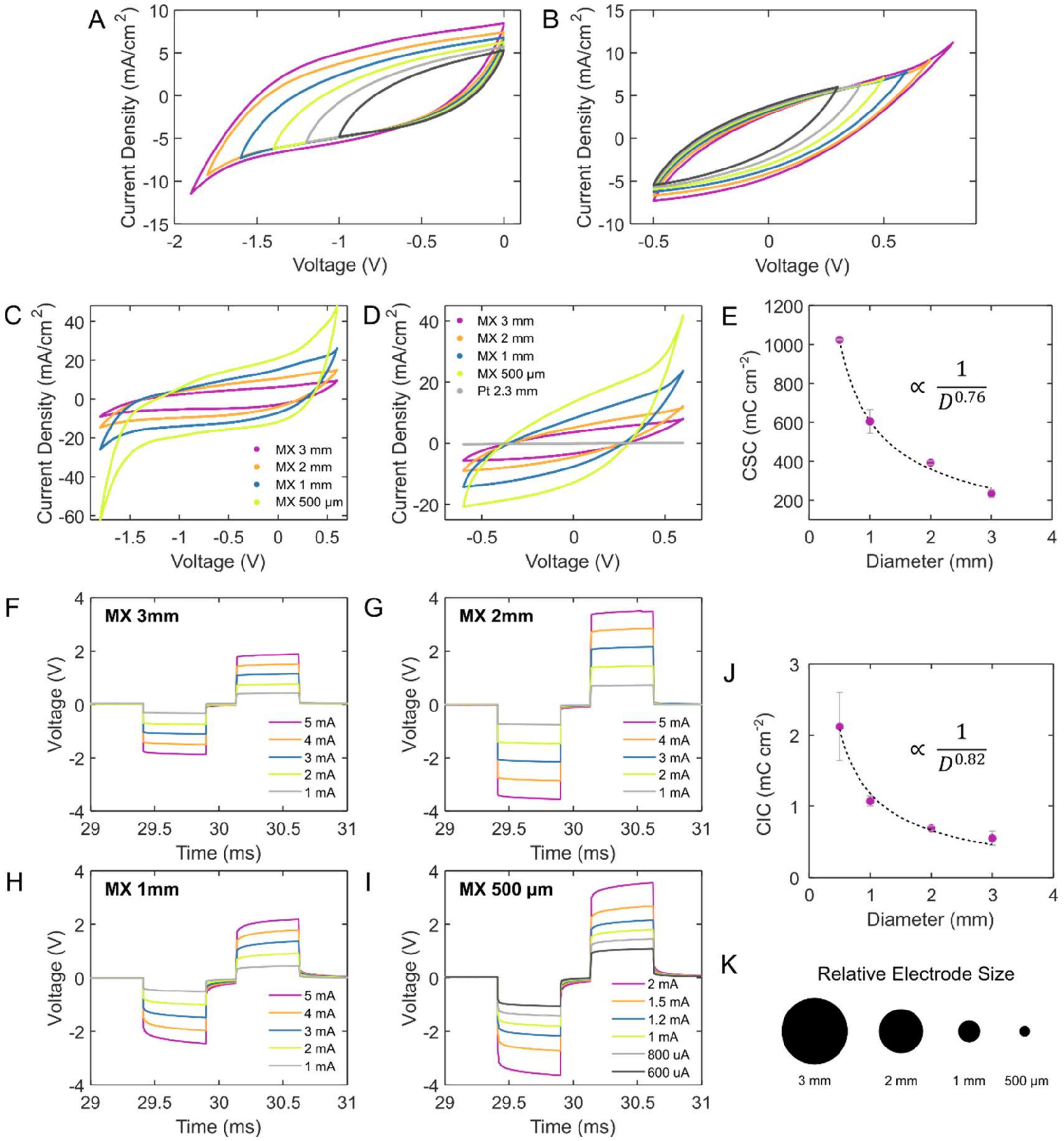


**Fig. S4. Scaling of electrochemical properties for MXtrodes.** (**A**) CVs probing the negative voltage limit for MXene at 50 mV/s. Hydrolysis of water begins at −1.9 V. (**B**) CVs probing the positive voltage limit for MXene. Current loss and evidence of faradaic current begins at +0.7 V. Test electrode for (A) and (B) was 3 mm-diameter planar MXene. (**C**) CVs in MXene safe voltage window, −1.8– +0.6 V, for 3 mm, 2 mm, 1 mm, and 500 µm-diameter MXtrodes. (**D**) CVs in MXene-Pt intersection window, −0.6 to +0.6 V, for 3 mm, 2 mm, 1 mm, and 500 µm-diameter planar MXtrodes and 2.3 mm-diameter Pt electrode. (**E**) Charge storage capacity of scaled planar MXtrodes as a function of diameter, highlighting the CSC scaling dependence on the electrode diameter, due to edge effects. CSC values were calculated for CVs in MXene water window. (**F-I**) Voltage transients for biphasic current pulses, with $t_c$ = $t_a$ = 500 µs and $t_{ip}$ = 250 µs, for currents ranging from 1 to 5 mA for (**F**) 3 mm, (**G**) 2 mm, (**H**) 1 mm, and (**I**) 500 µm-diameter planar MXene electrodes. (**J**) Charge injection capacity of scaled planar MXtrodes as a function of diameter, highlighting the CIC scaling dependence on the electrode diameter, due to edge effects. (**K**) Schematic demonstrating the relative electrode sizes used in the study.

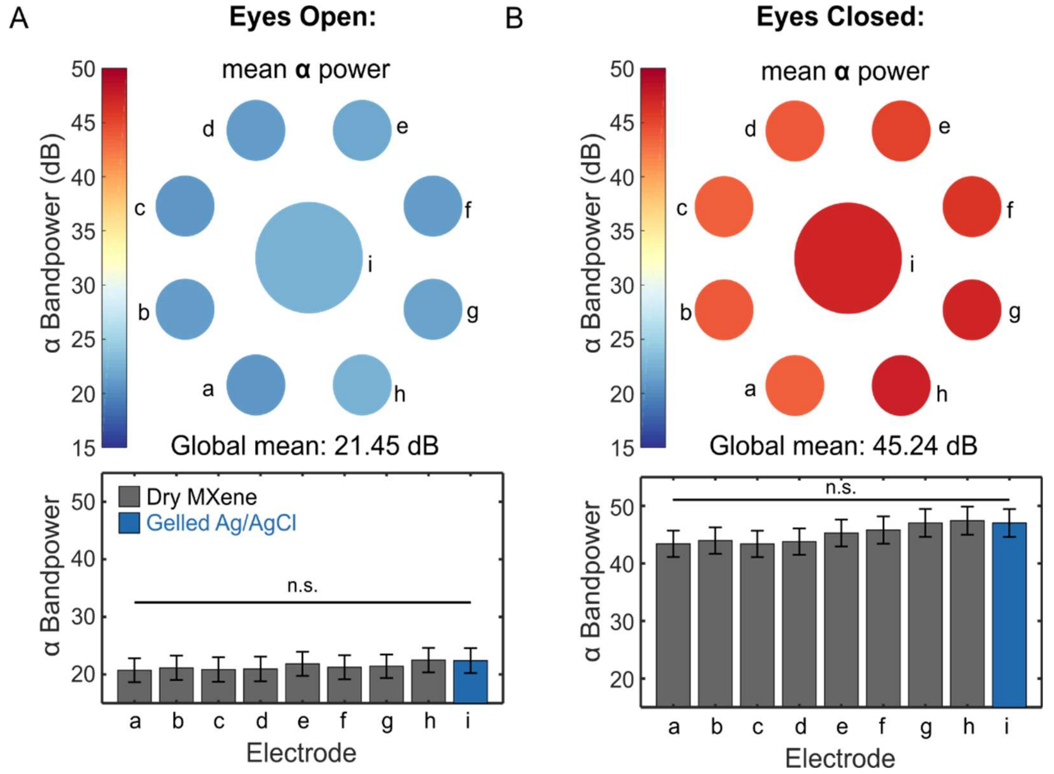


**Fig. S5. EEG alpha bandpower mapping.** (**A**) 8-12 Hz alpha bandpower across the 2 min recording in the eyes open state. Color plot (top) shows average alpha power for each electrode, mapped to its corresponding location on the scalp. Bar plot (bottom) shows average alpha power, with error bars corresponding to standard error of the mean across all time windows. No significant difference was detected between the gelled Ag/AgCl electrode (labelled i) and the dry MXtrode electrodes (labelled a-h). (**B**) The same alpha bandpower analysis shown in (A), for the eyes closed task.

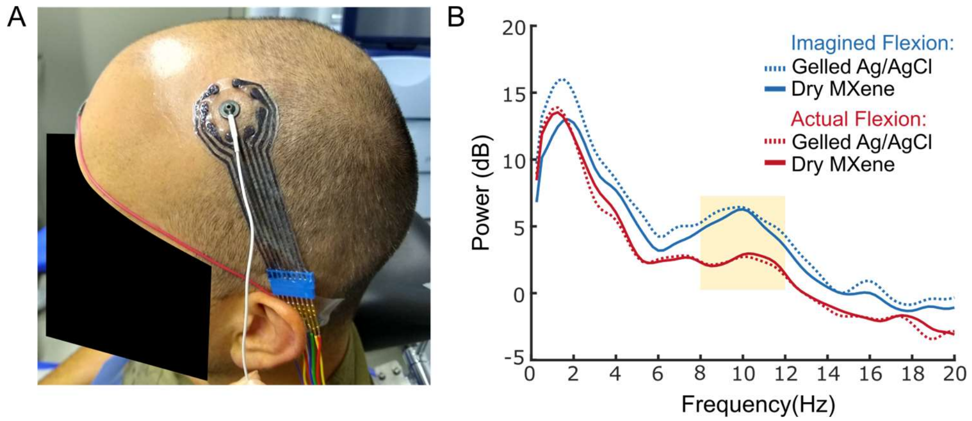


**Fig. S6. Motor EEG recording.** (**A**) Photograph of the EEG recording setup with electrodes centered over the hand motor region, as localized with single TMS pulses. (**B**) PSDs of the recorded EEG signal reveal a suppression of the 8-12 Hz motor mu rhythm during actual hand flexion as compared to imagined hand flexion.

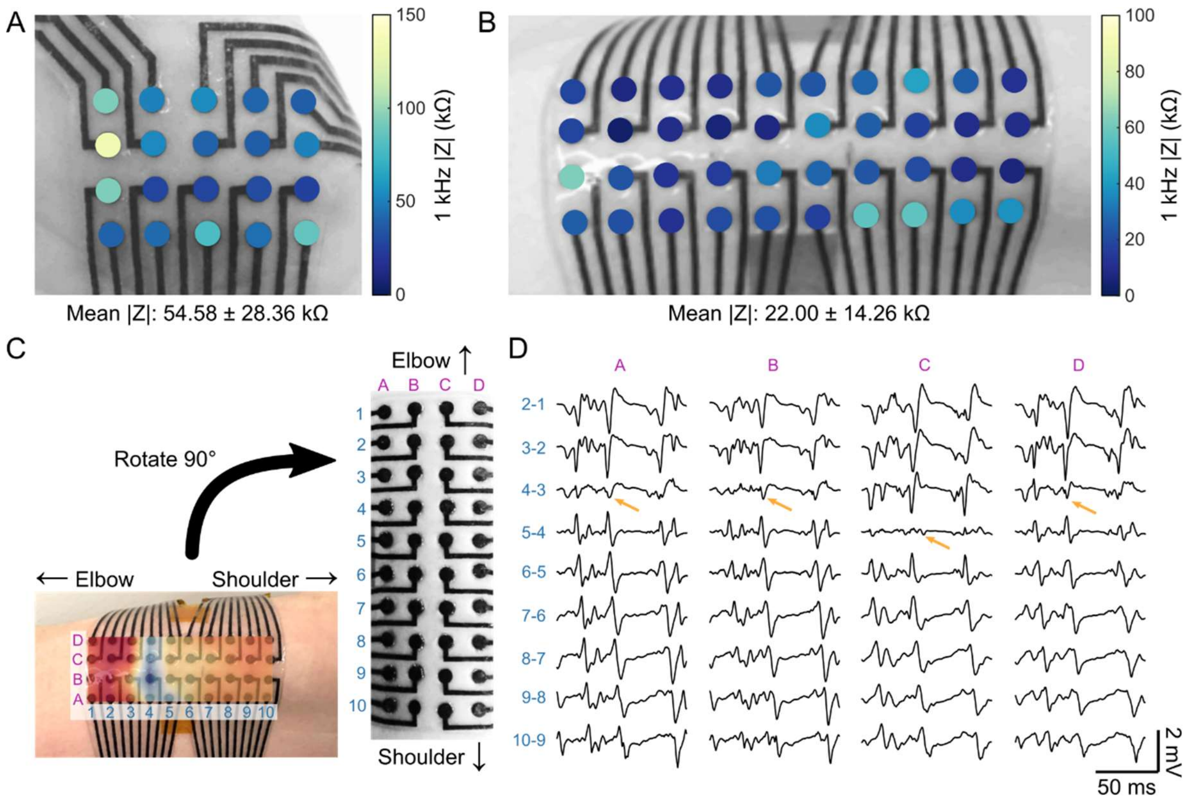


**Fig. S7. EMG array impedance and bipolar subtraction experiment.** (**A, B**) 1 kHz impedance magnitude maps for the (A) 20-ch planar MXtrode array used to map the APB muscle and the (B) 40-ch planar MXtrode array used to map the biceps, overlaid on images of the arrays on the subject during the experiment. (**C**) Schematic showing the arrangement for bipolar signal subtraction shown in (**D**) for resisted flexion EMG recordings on the biceps. The latency map obtained from the supraclavicular stimulation experiment is shown overlaid on the bottom image. (**D**) Bipolar EMG signals recorded during resisted flexion on the biceps. The location of the innervation zone is indicated by the arrows, and is clear from the propagation of the EMG signal outward from this region with some delay, as well as the inversion of the spikes on either side. The innervation zone determined from this analysis is in agreement with the region identified by the electrical stimulation-derived latency map shown in (C).

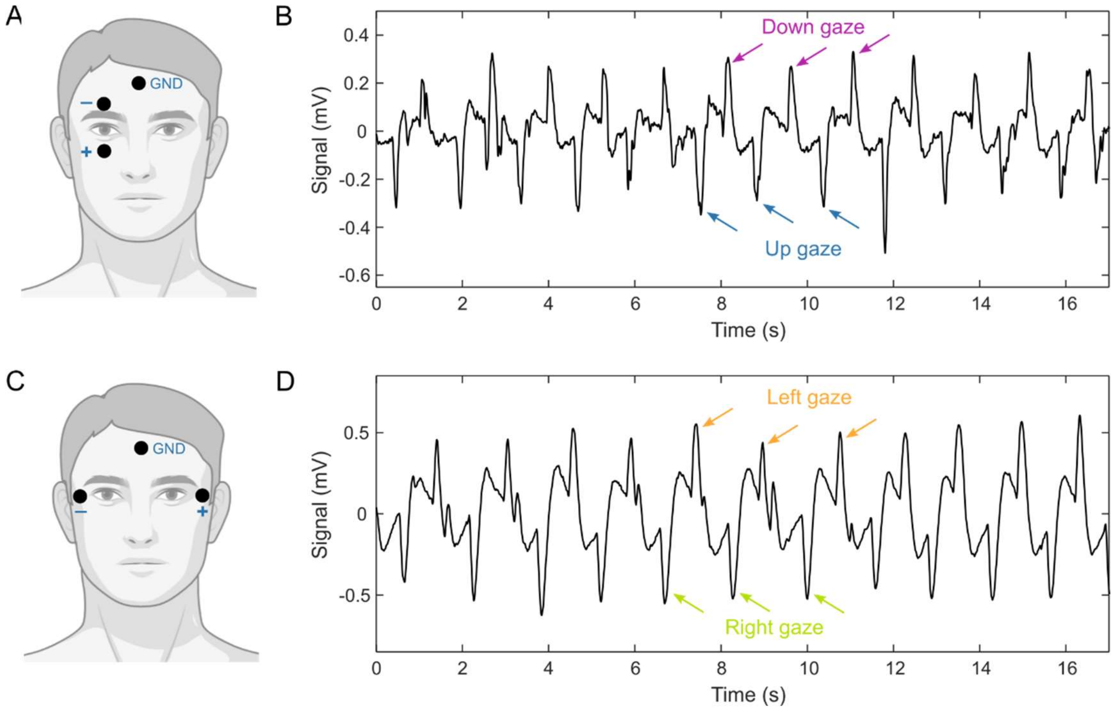


**Fig. S8. Electrooculography with MXtrodes.** (**A**) Schematic of EOG recording for monitoring up-down eye movements. (**B**) EOG data recorded on MXtrodes, showing distinct up and down eye movements. (**C**) Schematic of EOG recording for monitoring left-right eye movements. (**D**) EOG data recorded on MXtrodes, showing distinct left and right eye movements.

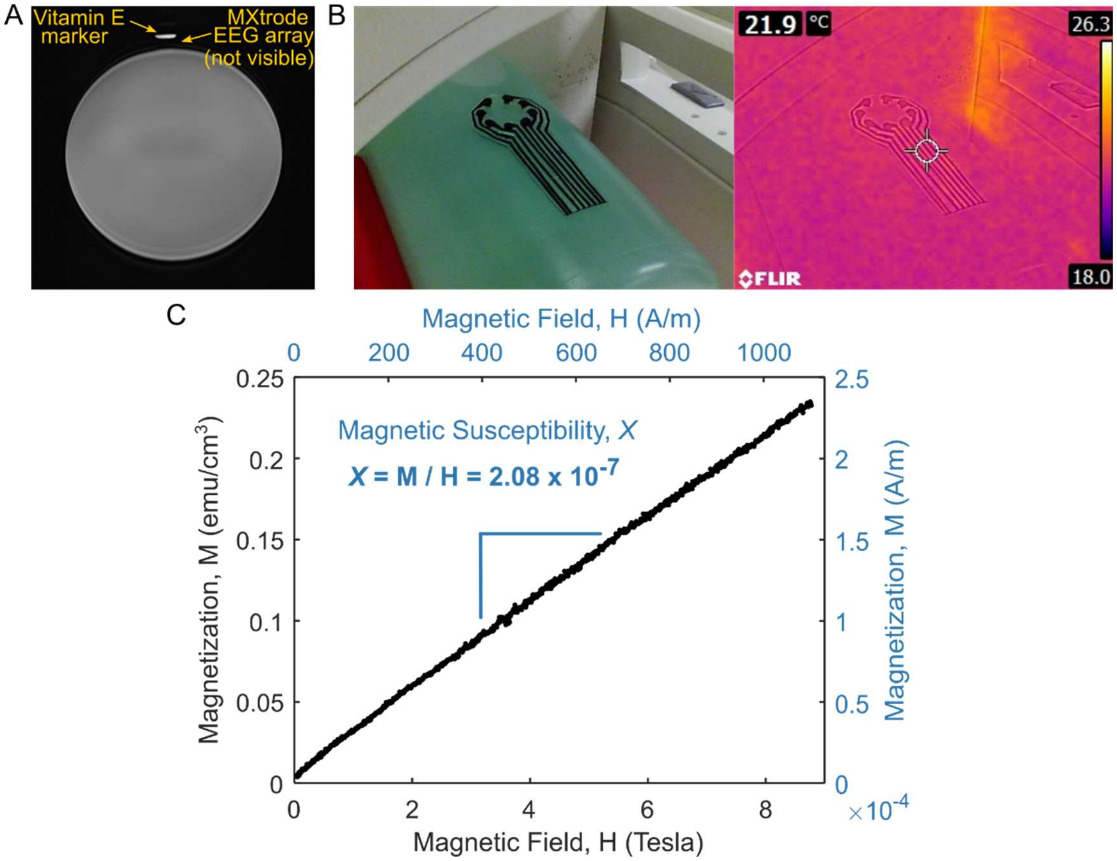


**Fig. S9. 3T MRI compatibility and magnetic susceptibility of MXene.** (**A**) MXtrode EEG array, imaged in a 3T clinical MRI with a T2 weighted sequence. The array was placed atop an MRI phantom and a Vitamin E marker was placed on top of the MXtrode array. The Vitamin E marker is visible, while the MXtrode array is not. (**B**) Thermal IR image of MXtrode EEG array captured immediately after a 10 min MRI sequence, showing no sign of heating. Left image shows MXtrode array atop the MRI phantom, and right image shows the thermal image overlay. (**C**) Magnetic susceptibility of $Ti_3C_2$ MXene measured at body temperature (i.e., 310 K) for applied magnetic field up to 9T. For calculation of the magnetic susceptibility value, *X*, magnetization and field strength were both converted to units of A/m such that *X* is unitless.

### Supplementary Tables

**Table S1. Electrochemical properties of MXtrode planar electrodes of varying diameters, compared to 2.3 mm-diameter Pt electrodes and to literature values for other electrode materials.** Scaling trends for each property are also shown in the top row.

| Material | Geometric Surface Area ($mm^2$) | 1 kHz \|Z\| in PBS (Ω)<br>↑ elec size = ↓ \|Z\| | Potential Limits vs. Ag/AgCl (V) | $CSC_C$ ($mC\ cm^{-2}$)<br>↑ elec size = ↓ CSC | $CIC_C$ ($mC\ cm^{-2}$)<br>↑ elec size = ↓ CIC | Source |
|---|---|---|---|---|---|---|
| MXene | 7.069 | 205.6 ± 11.1 | -1.8 – 0.6 | 233.1 ± 19.6 | 0.55 ± 0.10 | This work |
| | 3.142 | 369.2 ± 40.2 | -1.8 – 0.6 | 392.3 ± 9.8 | 0.69 ± 0.03 | This work |
| | 0.7854 | 430.2 ± 86.1 | -1.8 – 0.6 | 604.8 ± 61.4 | 1.07 ± 0.07 | This work |
| | 0.1963 | 729.2 ± 71.2 | -1.8 – 0.6 | 1024.1 ± 3.0 | 2.12 ± 0.48 | This work |
| Pt | 4.155 | 287.3 ± 7.8 | -0.6 – 0.8 | 4.3 ± 0.2 | 0.07 ± 0.003 | This work |
| Pt | 3.142 | 1910 [a] | -0.6 – 0.8 | 3.6 [a] | 0.17 [a] | *(59, 103)* |
| Au | 3.142 | 470 [a] | -0.9 – 0.6 | 0.3 [a] | 0.21 [a] | *(59, 103)* |
| PEDOT:PSS on Pt | 3.142 | 1690 [a] | -0.9 – 0.6 | 2.3 [a] | 0.18 [a] | *(59, 103)* |
| PEDOT:PSS on Au | 3.142 | 430 [a] | -0.9 – 0.6 | 1.0 [a] | 0.86 [a] | *(59, 103)* |
| Pt | 0.1963 | 7790 [a] | -0.6 – 0.8 | 5.1 [a] | 0.50 [a] | *(59, 103)* |
| Au | 0.1963 | 4430 [a] | -0.9 – 0.6 | 0.3 [a] | 0.21 [a] | *(59, 103)* |
| PEDOT:PSS on Pt | 0.1963 | 2110 [a] | -0.9 – 0.6 | 3.7 [a] | 0.66 [a] | *(59, 103)* |
| PEDOT:PSS on Au | 0.1963 | 990 [a] | -0.9 – 0.6 | 1.5 [a] | 1.35 [a] | *(59, 103)* |
| laser-roughened Pt | 0.7854 | 760 [a] | -0.6 – 0.8 | 2.1 ± 0.1 | 0.026 [a] | *(104)* |
| PEDOT/CNT coating on Pt | 0.0314 | 2100 [a] | -0.6 – 0.7 | 70 | 2.5 ± 0.1 [a] | *(105)* |
| Porous graphene (doped) | 0.09 | 519 | -1.3 – 0.8 | 50 | 3.1 | *(106)* |
| CNT fiber | 0.00145 | 11.2 ± 7.6 x$10^3$ | -1.5 – 1.5 | 372 ± 56 [b] | 6.52 | *(107)* |
| TiN | 0.004 | 11.5 x$10^3$ [a] | -0.9 – 0.9 | 2.47 | 0.55 | *(108)* |
| IrOx | 0.004 | 7.1 x$10^3$ [a] | -0.6 – 0.8 | 11 | 4 | *(108)* |
| PtIr | 5.985 | 125 | -0.7 – 0.7 | 5.0 | – | *(109)* |

[a] Represents values extracted from plots from the respective references.

[b] CSC calculated from CV in Pt window, -0.6 – 0.8 V at 100 mV/s.

**Table S2. $CSC_C$ values for planar MXtrodes of varying diameters and the comparison clinical Pt ECoG electrode.** CV scans were performed at 50 mV/s for each electrode in both its safe voltage window and the intersection of the MXene and Pt voltage windows.

| Material | Electrode diameter | $CSC_C$ (mC $cm^{-2}$) MXene window: -1.8 – 0.6 V | $CSC_C$ (mC $cm^{-2}$) Intersection: -0.6 – 0.6 V | $CSC_C$ (mC $cm^{-2}$) Pt window: -0.6 – 0.8 V |
|---|---|---|---|---|
| MXene | 3 mm | 233.1 ± 19.6 | 80.9 ± 8.9 | N/A |
| | 2 mm | 392.3 ± 9.8 | 118.6 ± 10.1 | N/A |
| | 1 mm | 604.8 ± 61.4 | 188.5 ± 28.4 | N/A |
| | 500 µm | 1024.1 ± 3.0 | 288.7 ± 1.7 | N/A |
| Pt | 2.3 mm | N/A | 4.0 ± 0.2 | 4.3 ± 0.2 |

### Supplementary Movies

**Movie S1. High-density dry EEG shows spatial patterns of alpha activation.** Alpha bandpower, calculated in 1 s windows with 0.5 s of overlap, mapped over the electrode locations. The outer ring of electrodes are the dry MXtrodes and the center electrode is the standard gelled Ag/AgCl electrode.

**Movie S2. Cortical stimulation with MXtrodes evokes whisker movement in rat.** Microstimulation of motor cortex with MXtrodes at 1.4 mV evokes whisker movements in anesthetized rat.
